# An optimized workflow to generate and characterize iPSC-derived motor neuron (MN) spheroids

**DOI:** 10.1101/2022.09.22.509079

**Authors:** Maria Jose Castellanos-Montiel, Mathilde Chaineau, Anna Kristyna Franco-Flores, Ghazal Haghi, Dulce Carrillo-Valenzuela, Wolfgang E. Reintsch, Carol X-Q Chen, Thomas M. Durcan

## Abstract

Motor neuron diseases (MNDs) are characterized by the progressive degeneration of motor neurons (MNs) from the cortex, brainstem and/or the spinal cord. In an effort to understand the underlying causes of this selective degeneration, a multitude of *in vitro* models based on induced pluripotent stem cell (iPSC)-derived MNs have been developed. Moreover, different groups have started to use advanced 3D structures, composed of MNs and other cell types to increase the physiological relevance of such *in vitro* models. For instance, spheroids are simple 3D models that have the potential to be generated in large numbers that can be used across different assays. In this study, we generated MN spheroids and developed a workflow to analyze them. We confirmed the expression of different MN markers as the MN spheroids differentiate, at both the transcript and protein level, as well as their capacity to display functional activity in the form of action potentials (APs) and bursts. We also identified the presence of other cell types, namely interneurons and oligodendrocytes, which share the same neural progenitor pool with MNs. In summary, we successfully developed a MN 3D model, and we optimized a workflow that can be applied to their characterization and analysis. In the future, we will apply this model and workflow to the study of MNDs by generating MN spheroids from patient-derived iPSC lines, aiming to contribute to the development of more advance and physiological *in vitro* disease models.

## 1 Introduction

Motor neurons (MNs) are a subset of efferent neurons within the nervous system responsible for innervating the muscles of the body, promoting their contraction through highly specialized and structurally organized synapses termed neuromuscular junctions (1). When the connection between MNs and skeletal muscle deteriorates or becomes interrupted as a result of MN degeneration, it leads to the development of a number of disorders known as motor neuron diseases (MNDs) which include amyotrophic lateral sclerosis (ALS), the most common disease in this category (2).

With the advent of induced pluripotent stem cell (iPSC) technology, new approaches have emerged to generate and culture different cell types of the human body *in vitro*, including MNs (3). Human iPSCs are generated by expressing the Yamanaka’s factors in adult somatic cells such as skin fibroblasts (FBs) or peripheral blood mononuclear cells (PBMCs) (4, 5). Thus, iPSCs can be derived from healthy individuals or patients with MNDs. In a disease context, patient-specific iPSCs retain the genetic background of the patient and when differentiated into MNs, they can display characteristics of the diseases *in vitro (6, 7)*. Remarkably, in cases where a monogenic mutation is identified as the primary cause of the disease, CRISPR-Cas9 genome editing, can be used to correct the mutation, generating isogenic controls that facilitate precise genotype and phenotype correlations (8–11). In addition to the latter advantage, iPSC-derived neurons allow researchers to study sporadic MND cases (6), which has never been achieved with other model systems. Taken together, these advantages open up new insights and therapeutic avenues to explore.

Most protocols available to generate iPSC-derived MNs are typically optimized to obtain two dimensional (2D) monolayer cultures (12–17) which have allowed the study of the pathogenic molecular mechanisms associated with MNDs such as cytoskeletal abnormalities, axonal transport deficits and changes in excitability (18, 19). However, several lines of evidence suggest that the absence of three dimensional (3D) cell-to-cell and cell-to-matrix interactions may be detrimental to the maturation (20, 21), differentiation (22) and morphology (23, 24) of cells grown *in vitro*. In addition, technical challenges arise when culturing iPSC-derived MNs as 2D monolayers. The cell bodies of the MNs often form clusters with their axons entangled (**Fig 1A**), making them inaccessible for some analyses. For instance, monitoring cell morphology, electrical activity and protein expression through immunocytochemistry becomes a challenging task (25, 26). Moreover, the clustering cells are prone to detachment from the surface, preventing their long-term culture. Maintaining cultures over several weeks is a critical aspect in MND modelling considering that disease phenotypes might only appear after a long period of culture. While some groups are optimizing 2D protocols to reduce the aggregation of iPSC-derived MNs (25, 27), other groups are developing techniques to generate 3D cultures which include spinal organoids (28–31) and MN spheroids (32–36) to address some of the technical issues that arise with 2D monolayers and to enhance MN maturation by exposing the cells to a 3D environment.

**Figure 1.**
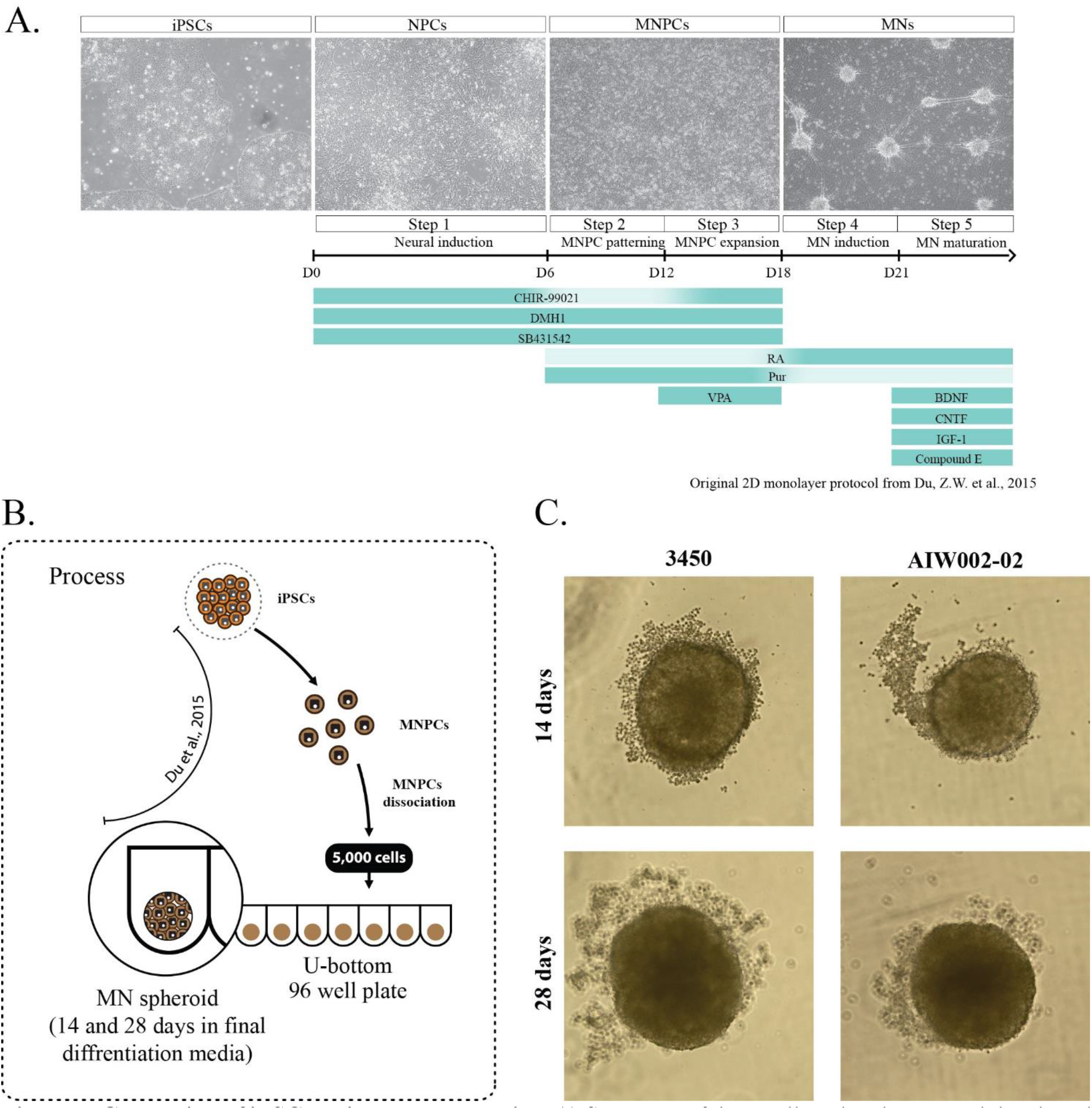
Generation of iPSC-derived MN spheroids. **A)** Summary of the small-molecule protocol developed by (13)and optimized by our group to generate iPSC-derived MNs as a 2D monolayer. Briefly, iPSCs are differentiated towards neural progenitor cells (NPCs) that are patterned to motor neuron neural progenitor cells (MNPCs) which are finally differentiated into MNs. **B)** At the MNPCs stage, two control cell lines (3450 and AIW002-02) were dissociated and plated into 96 U-bottom ultra-low attachment well plates in MN induction and maturation medium to generate MN spheroids that were kept in culture until their analysis. **C)** Representative bright-field pictures of MN spheroids after 14 and 28 days of differentiation of both control cell lines.

MN spheroids can be generated by making modifications to previously established 2D protocols (32–36). Plating motor neuron progenitor cells (MNPCs) onto a low attachment surface takes advantage of their ability to spontaneous assemble into clusters and form spheroids in which they can continue their differentiation towards MNs. Even though these MN spheroids show expression of well-known MN markers [i.e., pan-MN marker motor neuron and pancreas homeobox 1/ISL LIM homeobox 1 (HB9/ISL1) and choline acetyltransferase (CHAT)], a characterization using different markers to identify the presence of other cell types such as interneurons, oligodendrocytes and glial cells within MN spheroids is lacking. Additionally, the development and optimization of a defined workflow to analyze these 3D structures is a challenging task that needs to be addressed. With this in mind, we adapted a previously established 2D protocol (37) to generate 3D MN spheroids from iPSC-derived MNPCs that can be grown and maintained for up to 28 days, and longer if needed. By applying advanced 3D optical imaging and quantitative real-time PCR (qPCR) we characterized the identity of the cells within the MN spheroids and found that the majority of the cells expressed MN markers. Moreover, by performing microelectrode array (MEA) activity recordings we confirmed that these MN spheroids contain electrically active neurons. Easy to maintain and generated at high numbers, these spheres represent a simple model that has the potential to be applied to different assays and to be used as a cellular jigsaw, in which other cell types can be added to form more advanced co-culture spheroids.

## 2 Materials and equipment

### 2.1 Generation and maintenance of human iPSCs

Two control iPSC lines were used to optimize the protocols constituting the workflow for this study: AIW002-02 and 3450. The complete profiles of the iPSCs, culture conditions, and quality control analysis has been published (38). The 3450 and AIW002-02 iPSCs were maintained in E8 or mTeSR1 medium, respectively. Prior to the experiments, the iPSCs were found to be free from mycoplasma, hepatitis B/C and HIV 1/2 virus. The use of iPSCs in this research was approved by the McGill University Health Centre Research Ethics Board (DURCAN_IPSC / 2019-5374).

### 2.2 Generation of iPSC-derived MN spheroids

### 2.3 Cell Profiler macro for size profiling of MN spheroids

### 2.4 qPCR analysis of MN spheroids

### 2.5 Fixation, tissue clearing and immunofluorescent staining of MN spheroids

### 2.6 Microelectrode array (MEA) recordings of MN spheroids

## 3 Methods

### 3.1 Generation of iPSC-derived MN spheroids

Starting from iPSCs, we adapted a previously described protocol (13) to derive MNPCs (**Supplementary Figure 1**) that can be used to generate MN spheroids. The media and biochemicals used to generate MN spheroids are listed in **Table 1**. Consumables and equipment are listed in **Table 2**. The media compositions for the different differentiation steps are listed in **Table 3**.

**Table 1.**
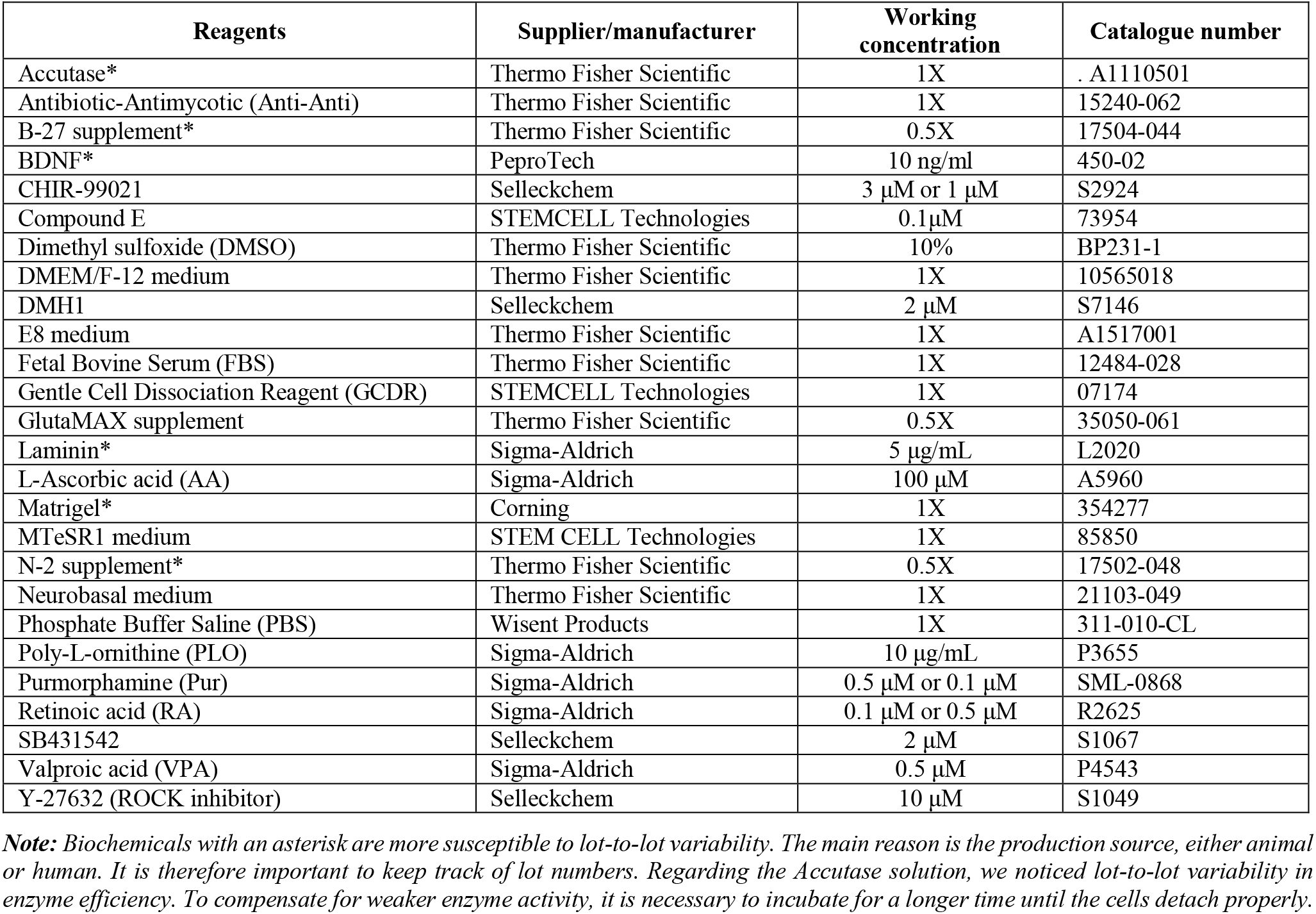
List of media and biochemicals.

**Table 2.** List of consumables and equipment.

| Consumables and equipment | Supplier | Catalogue number |
| --- | --- | --- |
| 100 mm culture dishes | Thermo Fisher Scientific | 353003 |
| 96 round bottom ultra-low attachment plates | Corning | CLS7007 |
| Cell culture incubator | Thermo Fisher Scientific | Steri-Cycle Model 370 Ref#20 |
| Cell scraper | Corning | CLS3010 |
| Centrifuge | Eppendorf | 022626001 |
| Centrifuge with adapter for culture well plates | Eppendorf | 5810 |
| Conical tube, 15mL | Thermo Fisher Scientific | 352097 |
| Cryovials | Sarstedt | 72.379 |
| Luna-II™ automated cell counter | Logos biosystems | L40002 |
| Plastic serological pipet, 10mL | Thermo Fisher Scientific | 13-678-11E |
| Plastic serological pipet, 1mL | Thermo Fisher Scientific | 13-678-11B |
| Plastic serological pipet, 5mL | Thermo Fisher Scientific | 13-678-11D |
| T-25 flasks | Thermo Fisher Scientific | 12-556-009 |
| T-75 flasks | Thermo Fisher Scientific | 12-556-010 |

**Table 3.**
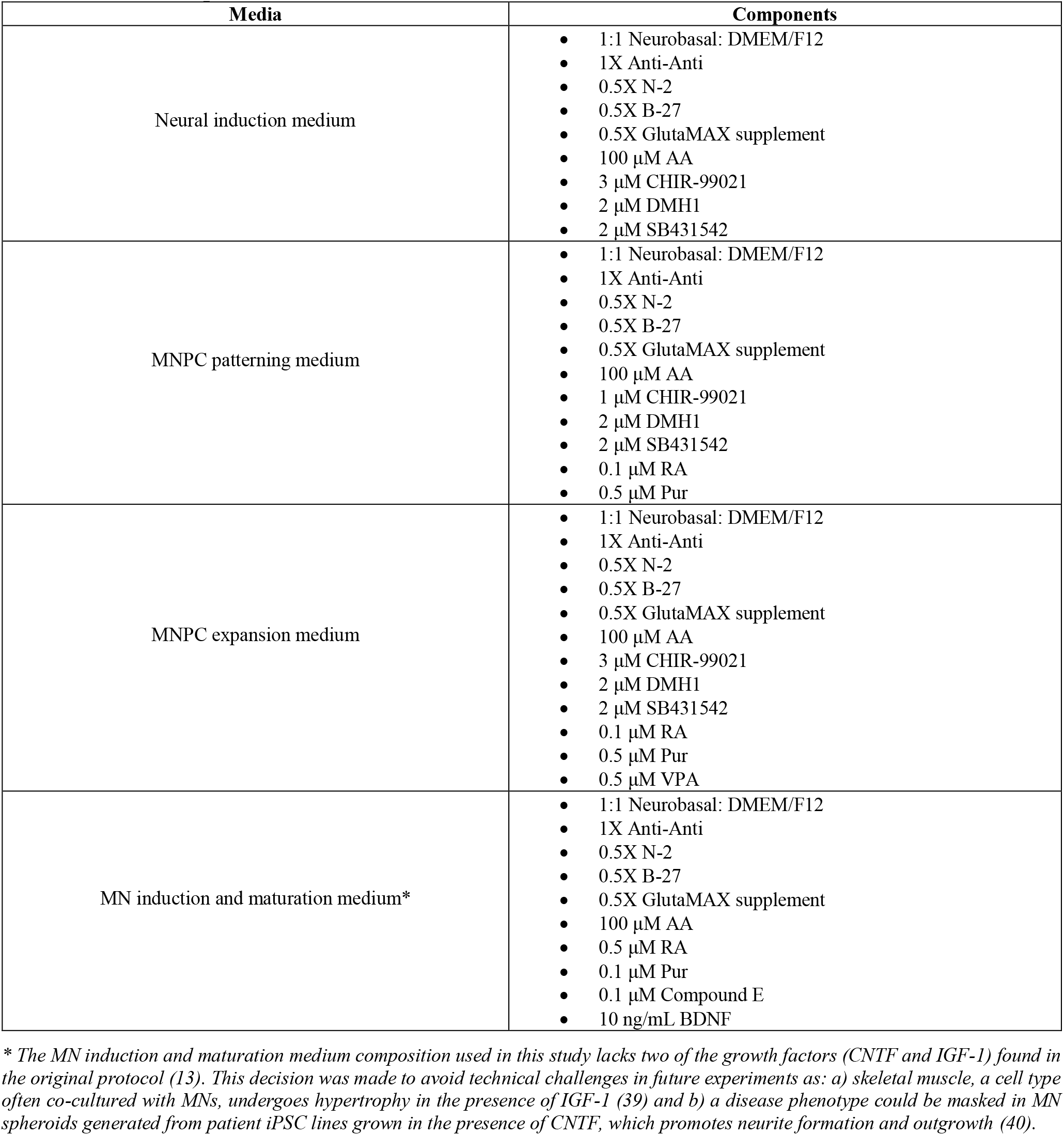
Media composition.

| Media | Components |
| --- | --- |
| Neural induction medium | • 1:1 Neurobasal: DMEM/F12<br>• 1X Anti-Anti<br>• 0.5X N-2<br>• 0.5X B-27<br>• 0.5X GlutaMAX supplement<br>• 100 μM AA<br>• 3 μM CHIR-99021<br>• 2 μM DMH1<br>• 2 μM SB431542 |
| MNPC patterning medium | • 1:1 Neurobasal: DMEM/F12<br>• 1X Anti-Anti<br>• 0.5X N-2<br>• 0.5X B-27<br>• 0.5X GlutaMAX supplement<br>• 100 μM AA<br>• 1 μM CHIR-99021<br>• 2 μM DMH1<br>• 2 μM SB431542<br>• 0.1 μM RA<br>• 0.5 μM Pur |
| MNPC expansion medium | • 1:1 Neurobasal: DMEM/F12<br>• 1X Anti-Anti<br>• 0.5X N-2<br>• 0.5X B-27<br>• 0.5X GlutaMAX supplement<br>• 100 μM AA<br>• 3 μM CHIR-99021<br>• 2 μM DMH1<br>• 2 μM SB431542<br>• 0.1 μM RA<br>• 0.5 μM Pur<br>• 0.5 μM VPA |
| MN induction and maturation medium* | • 1:1 Neurobasal: DMEM/F12<br>• 1X Anti-Anti<br>• 0.5X N-2<br>• 0.5X B-27<br>• 0.5X GlutaMAX supplement<br>• 100 μM AA<br>• 0.5 μM RA<br>• 0.1 μM Pur<br>• 0.1 μM Compound E<br>• 10 ng/mL BDNF |

\* The MN induction and maturation medium composition used in this study lacks two of the growth factors (CNTF and IGF-1) found in the original protocol (13). This decision was made to avoid technical challenges in future experiments as: a) skeletal muscle, a cell type often co-cultured with MNs, undergoes hypertrophy in the presence of IGF-1 (39) and b) a disease phenotype could be masked in MN spheroids generated from patient iPSC lines grown in the presence of CNTF, which promotes neurite formation and outgrowth (40).

#### 3.1.1 Day 0 (D0): Seeding of iPSCs

Start from a 10 mm dish of human iPSCs with a confluence between 70-80%, containing less than 5% of spontaneously differentiated cells. Once cells are ready, aspirate the medium from the iPSC culture and wash with 5 mL of DMEM/F-12 containing 1X Antibiotic-Antimycotic (Anti-Anti). Add 5mL of Gentle Cell Dissociation Reagent (GCDR) and incubate at room temperature (RT) for 4-5 min. Avoid incubating the iPSCs in GCDR for too long as cells dissociated into single cells are not ideal for the induction process. Aspirate the GCDR and rinse the cells with 5 mL of DMEM/F-12 containing 1X Anti-Anti. Aspirate the medium and then add 5 mL of DMEM/F-12 containing 1X Anti-Anti. Gently detach the colonies with a cell scraper and transfer the cells to a 15 mL conical tube. Wash the dish with an additional 5 mL of DMEM/F-12 containing 1X Anti-Anti to collect the remaining detached cells. Determine the cell number using a cell counter. Transfer 2-3 million cells to a fresh 15 mL conical tube and pellet the cells by centrifugation at 1200 rpm for 3 min. Aspirate the supernatant. Gently resuspend the cells in 1 mL of neural induction medium containing ROCK inhibitor (10 μM). Transfer the cells to a Matrigel-coated T-25 flask and add 4 mL of neural induction medium containing ROCK inhibitor (10 μM). Distribute the cells uniformly and place the flask in a 37°C/5%CO_2_ incubator.

The quality of the iPSCs (38) is crucial to the successful generation of MNPCs. Thus, mycoplasma tests should be performed routinely, ideally every other week to confirm the absence of contamination before using the iPSCs. To verify the absence of mycoplasma from cultures, we used a mycoplasma detection assay (MycoAlert™ mycoplasma detection kit, Lonza) to measure luminescence of 1.5 mL medium samples, according to manufacturer’s instructions.

#### 3.1.2 Day 1 (D1): Neural induction

Within 24 h of plating the iPSCs, change the medium to 5 mL of neural induction medium without ROCK inhibitor. Continue to change the medium every other day until D6 to induce the cells towards a neural progenitor cell (NPC) identity.

Check the cell morphology and density of the cells 24 h after plating. The cells must be 30% confluent. If the cells are too confluent causing the medium to turn yellow, change the medium every day until D6.

#### 3.1.3 Day 6 (D6): MNPC patterning

Six days after plating the iPSCs in neural induction medium, the cells should be 100% confluent and differentiated into NPCs. For the next step, cells will be plated on a T-25 and a T-75 PLO/laminin-coated flasks. For PLO/laminin coating, PLO must be kept for at least 2 h or overnight, then washed three times with 1X PBS and switched to laminin for a minimum of 2 h or overnight. Aspirate the neural induction medium and rinse the cells with 5 mL of DMEM/F-12 containing 1X Anti-Anti. Add 2 mL of GCDR and incubate at 37°C for 5-7 min. After the incubation, cells will begin lifting from the dish. Gently tap the plate to allow for the complete detachment of the cells into the GCDR. Add 5 mL of DMEM/F-12 containing 1X Anti-Anti and transfer the cells to a 15 mL conical tube. Pellet the cells by centrifugation at 1200 rpm for 3 min. Aspirate the supernatant. Resuspend the cells in 4 mL of MNPC patterning medium containing ROCK inhibitor (10 μM) and gently pipette the cells up and down. Plate 3 mL of the cell suspension into the PLO/laminin-coated T-75 flask. Complete with 12 mL of MNPC patterning medium containing ROCK inhibitor (10 μM) to reach a final volume of 15 mL and distribute the cells uniformly. Plate the remaining mL of the cell suspension into the PLO/laminin-coated T-25 flask. Complete with 4 mL of MNPC patterning medium containing ROCK inhibitor (10 μM) to reach a final volume of 5 mL and distribute the cells uniformly. Place the flasks in a 37°C/5%CO_2_ incubator and 24 h after plating the NPCs, change the medium to MNPC patterning without ROCK inhibitor. Continue to change the MNPC patterning medium every other day until D12.

Check the cell morphology and density of the cells 24 h after plating. The cells must be 50% confluent. If the cells are too confluent causing the medium to turn yellow, change the medium every day until D12.

#### 3.1.4 Day 12 (D12): MNPC expansion

Six days after plating the NPCs in MNPC patterning medium, cells should be 70% to 100% confluent and patterned to MNPCs. For the next step, cells will be plated into four T-75 PLO/laminin-coated flasks. Split the cells following section **3.1.3**. Resuspend the cells in 4 mL of MNPC expansion medium containing ROCK inhibitor (10 μM) and gently pipette the cells up and down. Plate 1 mL of cell suspension into each of the four PLO/laminin-coated T-75 flasks and distribute the cells uniformly. Complete with 14 mL of MNPC expansion medium containing ROCK inhibitor (10 μM) to reach a final volume of 15 mL and distribute the cells uniformly. Place the flasks in a 37°C/5% CO_2_ incubator. 24 h after plating the MNPCs, change the medium to MNPC expansion without ROCK inhibitor. Continue to change the medium every other day until D18.

If the cells are too confluent causing the medium to turn yellow, change the medium every day until D18.

#### 3.1.5 Day 18 (D18): MNPC storage, split or MN spheroid generation

##### MNPC storage

Six days after plating the MNPCs in MNPC expansion medium, cells should be 70% to 100% confluent. Proceed directly to freeze or split the cells. MNPCs can be frozen up to passage 3, depending on the cell morphology and density of the cells. MNPCs reduce their proliferation after four or more passages. The four T-75 flasks in MNPC expansion medium can be frozen into 20 to 40 cryovials, with each vial containing ~3-5 million cells in 1 mL of FBS containing 10% DMSO.

##### MNPC thawing

Thaw the frozen cryovial of MNPCs in a 37°C water bath by gently shaking the cryovial continuously until only a small, frozen cell pellet remains. Sterilize the outside of the cryovial with 70% ethanol. Transfer the cells to a 15 mL conical tube with 4 mL of DMEM/F-12 containing 1X Anti-Anti and pipette gently. Pellet the cells by centrifugation at 1200 rpm for 3 min. Aspirate the supernatant. Resuspend the cells in 5 mL of MNPC expansion medium containing ROCK inhibitor (10 μM) and transfer them into a T-25 PLO/laminin-coated flask. Place the flask in a 37°C/5% CO_2_ incubator. 24 h after plating the MNPCs, change the medium to MNPC expansion without ROCK inhibitor. Continue to change the MNPC expansion medium every other day for 6-7 days.

##### MNPC split

Before starting MN spheroid generation from a recently thawed MNPC cryovial, it is recommended to passage the cells at least once to allow complete recovery of the cells. After each passage, plate two T-25 flasks. One flask will be used to keep the MNPC stock in culture (up to 5 passages) through GCDR dissociation, and the other flask will be used to initiate MN spheroid generation through Accutase dissociation. Split the cells following section **3.1.3**. Resuspend the cells in MNPC expansion medium containing ROCK inhibitor (10 μM) and plate ~3-4 million cells into a PLO/laminin-coated T-25 flask. Complete with MNPC expansion medium containing ROCK inhibitor (10 μM) to reach a final volume of 5 mL, distribute the cells uniformly and place the flask in a 37°C/5% CO_2_ incubator. Within 24 h of plating the MNPCs, change the medium to MNPC expansion without ROCK inhibitor. Continue to change the MNPC expansion medium every other day until 6-7 days.

##### MN spheroid generation

MNPCs are ready for differentiation when cells have been cultured in MNPC expansion medium for 6-7 days. Split the cells following section **3.1.3**., incubating with 2 mL of Accutase at 37°C for 4-6 min instead of GCDR to achieve a single cell suspension. Monitor the cells so the dissociation can be stopped as soon as all cells have detached. Resuspend the cells in 1 mL of induction and maturation medium without ROCK inhibitor and pipette the cells up and down. Quantify cell number using a cell counter. Plate 5,000 MNPCs in 100 μl into each well of a 96 U-bottom ultra-low attachment plate in MN induction and maturation medium (**Fig 1B**). Centrifuge the plate at 1200 rpm for 5 min using a centrifuge with adapters for cell culture plates. Place the culture plate in a 37°C/5% CO_2_ incubator. This centrifugation step will speed up the aggregation of the cells, thus speeding up the formation of the spheroids. However, the spheroids will also form (albeit at a slower rate) even if this step is not performed. To maintain the MN spheroids in culture over time, replenish each well of the 96 U-bottom ultra-low attachment plate with 50 μl of MN induction and maturation medium every 14 days.

Note: When plating the cells into 96 U-bottom ultra-low attachment plate, avoid seeding cells in the outer wells of the plate. Evaporation of the medium is higher in these wells and it has a negative impact on spheroid formation. Instead, fill the outer wells with 200 μl of 1X PBS.

### 3.2 Cell Profiler macro for size profiling of MN spheroids

The equipment used to profile the size of the MN spheroids is listed in **Table 4**. Bright-field images of MN spheroids were acquired with a light microscope after culture for 14 and 28 days in MN induction and maturation medium (**Fig 1C**). Importantly, the images acquired at the two different time points were always saved with their location within the plate (Ex. B2, B3, B4, B5…) in order to follow each MN spheroid growth over time. We performed measurements on 28-30 MN spheroids from each of five different batches for each cell line (AIW002-02=150 spheroids; 3450=145 spheroids). The images were analyzed by a pipeline developed in our group using CellProfiler Analyst, www.cellprofiler.org (41), and available online (https://doi.org/10.17605/OSF.IO/V84WS). Briefly, images of the spheroids obtained with a bright-field microscope are inverted and speckles with a diameter smaller than 10 pixels are filtered out to eliminate cell debris and cells that were not incorporated into the spheroids. From this point, the primary object (named “Sphere”) is identified using Otsu thresholding and its size and shape are measured (**Supplementary Figure 2A**). The primary objects identified as the final spheroids are overlaid with the original input image for visual control of the spheroid identification (**Supplementary Figure 2B**). As a result, the pipeline gives a CSV file with different measurements in pixels for each MN spheroid including the min Feret diameter (used as diameter) and the area. The measurements in pixels have to be manually transformed into μm (diameter) or μm^2^ (area) according to the microscope scale if needed. The radius of each spheroid was calculated by dividing the min Feret diameter by half and substitute it into the formula (4/3∏r^3^) to determine the volume of a sphere (**Fig 2A/B**). In addition, to assess the circularity of the spheroids, the radius of each spheroid was substituted into the formula (∏r^2^) to determine the area of a circle and a ratio between the calculated area and the area given by the software (**Fig 2C**) was performed. The efficiency of the CellProfiler pipeline to identify the primary objects (spheroids) highly depends on the pixel intensities through the entire bright field image, meaning that sometimes cell debris or individual cells with a pixel intensity similar to the sphere lead to misinterpretation of the primary object (**Supplementary Figure 2C**). To establish a threshold that helped us define images in which the CellProfiler pipeline performed poorly, we used GraphPad Prism version 9.1.1 to identify outliers using the ROUT method with a Q=1%. The images marked as potential outliers were checked to corroborate they were not the actual size of the spheroid. A paired t-student was performed to compare the diameter, area and volume measurements to compare 14- and 28-days MN spheroids (AIW002-02=137 spheroids; 3450=142 spheroids).

**Table 4.** List of equipment.

| Equipment | Supplier | Catalogue number |
| --- | --- | --- |
| EVOS XL Core | Thermo Fisher Scientific | AMEX1000 |
| JuLI<sup>TM</sup> Stage | NanoEntek | NANOJS1000S |

**Figure 2.**
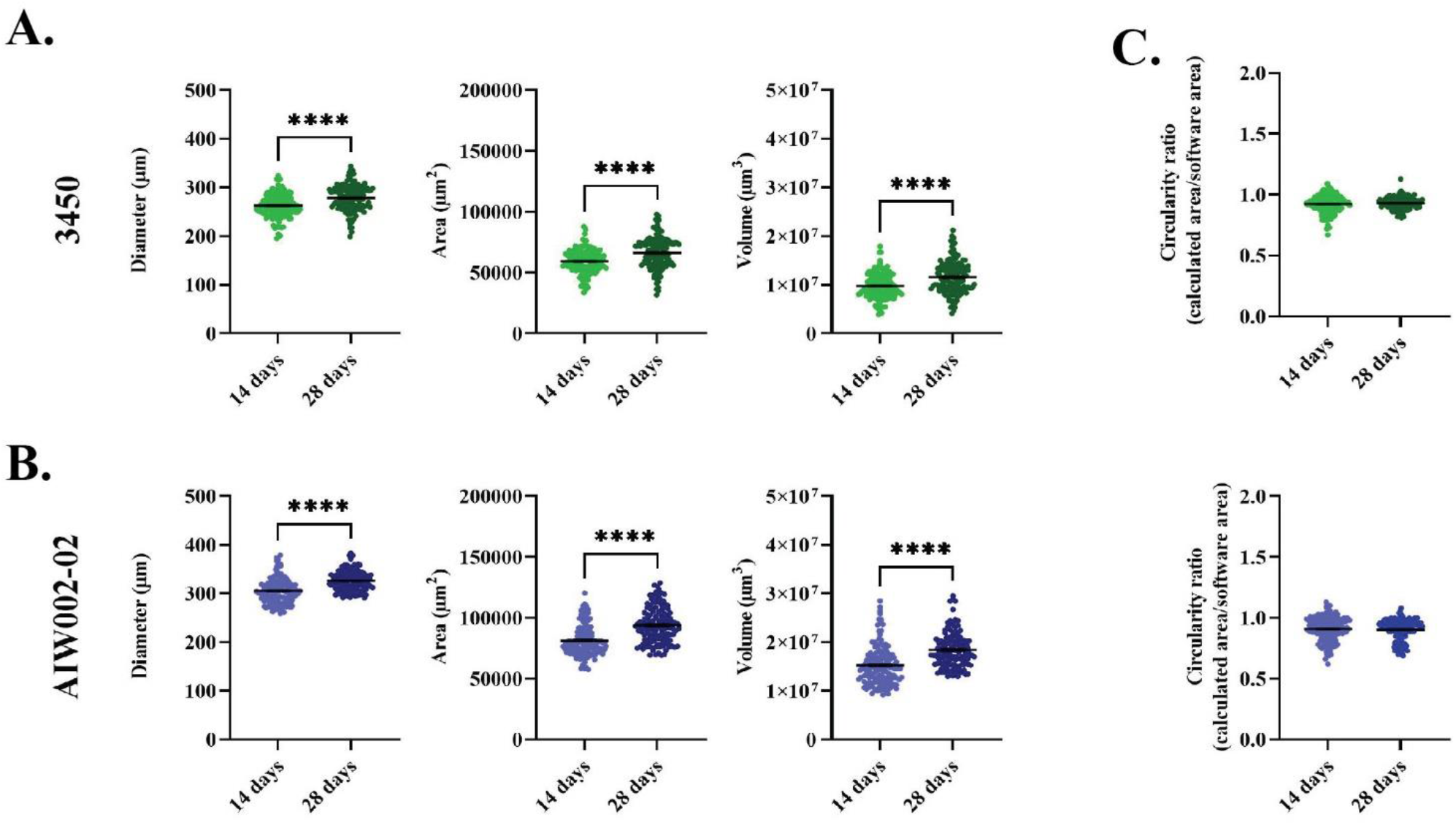
Size profile by a Cell Profiler pipeline. iPSC-derived MN spheroids from the **A)** 3450 and **B)** AIW002-02 control lines are consistently generated across different batches. They exhibit an increase in size over time as shown by diameter, area and volume measurements. Scatter plots show the mean ± SEM; for each cell line, five MN spheroid batches coming from at least two iPSC-derived MNPCs batches induced through independent differentiation processes (AIW002-02=137 spheroids; 3450=142 spheroids). To ensure that MNPC passage number was not having any effect on MN spheroid formation, we used the MNPCs at passage number 2 to 4. Significance was determined by a paired *t*-student. ****p<0.0001. **C)** Additionally, MN spheroids display a circular shape with bulges appearing occasionally but without disturbing the circular morphology as shown by the circularity ratio (0= not circular; 1=perfect circle).

### 3.3 qPCR analysis of MN spheroids

MN spheroids from one 96 U-bottom ultra-low attachment plate (60 spheroids) were pooled at 14 and 28 days for RNA extraction. The materials and reagents to perform the qPCR from MN spheroid samples are listed in **Table 5**. The consumables and equipment are listed in **Table 6**. The probes used for the qPCR analysis are listed in **Table 7**.

**Table 5.** List of material and reagents.

| Reagents | Supplier/manufacturer | Working concentration | Catalogue number |
| --- | --- | --- | --- |
| 2X no UNG Taqman™ fast advanced master mix | Thermo Fisher Scientific | N/A | A44360 |
| dNTPs | New England BioLabs | 0.5 mM | N0447L |
| DTT | Thermo Fisher Scientific | 0.01 M | 28025013 |
| First strand buffer | Thermo Fisher Scientific | 1X | 28025013 |
| miRNeasy micro kit | Qiagen | N/A | 217004 |
| M-MLV RT | Thermo Fisher Scientific | 400 U | 28025013 |
| Random primers | Thermo Fisher Scientific | 12.5 ng/μl | 48190011 |

**Table 6.** List of consumables and equipment.

| Consumables and equipment | Supplier | Catalogue number |
| --- | --- | --- |
| MicroAmp™ 8 tube strip (0.2 mL) | Thermo Fisher Scientific | N8010580 |
| MicroAmp™ optical 384-well plate reaction plate | Thermo Fisher Scientific | 4309849 |
| MicroAmp™ optical 8-cap strips | Thermo Fisher Scientific | 4323032 |
| MicroAmp™ optical adhesive | Thermo Fisher Scientific | 4311971 |
| Nanodrop | Thermo Fisher Scientific | ND-ONE-W |
| QuantStudio 5 | Thermo Fisher Scientific | A28140 |
| SimpliAmp™ thermal cycler | Thermo Fisher Scientific | A24811 |

**Table 7.**
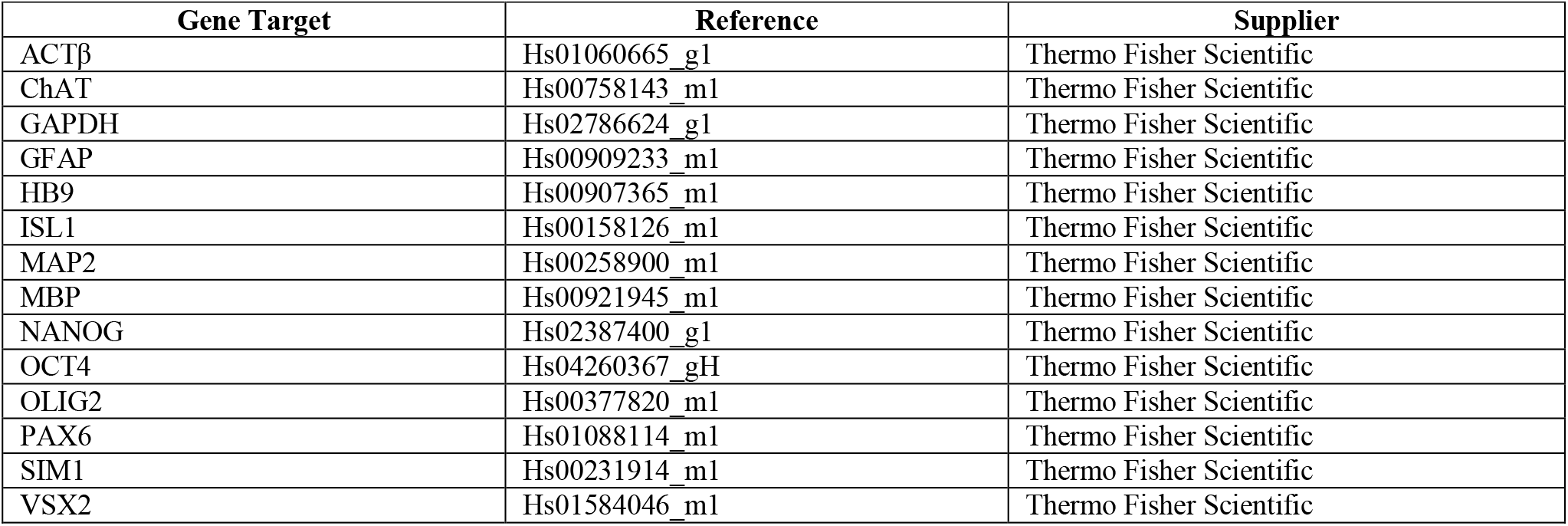
Probes/primers.

| Gene Target | Reference | Supplier |
| --- | --- | --- |
| ACTβ | Hs01060665_g1 | Thermo Fisher Scientific |
| ChAT | Hs00758143_m1 | Thermo Fisher Scientific |
| GAPDH | Hs02786624_g1 | Thermo Fisher Scientific |
| GFAP | Hs00909233_m1 | Thermo Fisher Scientific |
| HB9 | Hs00907365_m1 | Thermo Fisher Scientific |
| ISL1 | Hs00158126_m1 | Thermo Fisher Scientific |
| MAP2 | Hs00258900_m1 | Thermo Fisher Scientific |
| MBP | Hs00921945_m1 | Thermo Fisher Scientific |
| NANOG | Hs02387400_g1 | Thermo Fisher Scientific |
| OCT4 | Hs04260367_gH | Thermo Fisher Scientific |
| OLIG2 | Hs00377820_m1 | Thermo Fisher Scientific |
| PAX6 | Hs01088114_m1 | Thermo Fisher Scientific |
| SIM1 | Hs00231914_m1 | Thermo Fisher Scientific |
| VSX2 | Hs01584046_m1 | Thermo Fisher Scientific |

Total RNA was isolated using the miRNeasy micro kit according to manufacturer’s instructions. To obtain cDNA, reverse transcriptions were done by mixing 40 ng of total RNA, 12.5 ng/μl of random primers, 0.5 mM dNTPs, 0.01 M DTT, 1X first strand buffer and 400 U M-MLV RT in a total volume of 40 μl. qPCR reactions were performed in 384-well plates using the QuantStudio5 PCR machine. For each well, the PCR mix included 9 μl of 2X no UNG Taqman™ Fast Advanced Master, 0.5 μl of primers/probe mix, 1 μl of cDNA, with H_2_O up to 10 μl. Serial dilutions of a mix of cDNA, consisting of cDNA from all the samples, ranging between 50 ng and 0.003052 ng were used to generate a calibration curve for absolute quantification. Expression levels were given as a ratio between the relative quantities of the gene of interest and the endogenous control. The mean between Actβ and GAPDH was used as the endogenous controls for normalization.

### 3.4 Fixation, tissue clearing and immunofluorescent staining of MN spheroids

MN spheroids from 96 U-bottom ultra-low attachment plates (60 spheroids/plate) were fixed, cleared and immunostained at 14 and 28 days respectively. The reagents to perform the fixation and the immunofluorescent staining of the MN spheroids are listed in **Table 8**. The compositions to prepare the two solutions needed to perform the Clear Unobstructed Brain/Body Imaging Cocktail and Computational Analysis (CUBIC) protocol (42), CUBIC reagent 1 (R1) and CUBIC reagent 2 (R2), are listed in **Table 9** and **Table 10** respectively. The composition of the blocking solution used for the immunostaining process is listed in **Table 11**. The consumables and equipment are listed in **Table 12**. The primary and secondary antibodies used for immunostaining are listed in **Table 13** and **Table 14** respectively.

**Table 8.** List of material and reagents.

| Reagents | Supplier/manufacturer | Working concentration | Catalogue number |
| --- | --- | --- | --- |
| 16% formaldehyde (FA) | Thermo Fisher Scientific | 4% | 28908 |
| Hoechst33342 | Thermo Fisher Scientific | 1:1000 | H3570 |
| Phosphate Buffer Saline (PBS) | Wisent Bioproducts | 1X | 311-010-CL |

**Table 9.** Cubic reagent 1 (R1) composition.

| Reagents | Supplier/manufacturer | Working concentration | Catalogue number |
| --- | --- | --- | --- |
| dH<sub>2</sub>O | N/A | 35% by wt | N/A |
| Urea | Sigma-Aldrich | 25% by wt | 15604 |
| Quadrol (N, N, N', N'-tetrakis (2-hydroxy-propyl) ethylenediamine) | Sigma-Aldrich | 25% by wt | 122262 |
| Triton X-100 | Bioshop | 15% by wt | TRX506 |

**Table 10.** Cubic reagent 2 (R2) composition.

| Reagents | Supplier/manufacturer | Working concentration | Catalogue number |
| --- | --- | --- | --- |
| dH<sub>2</sub>O | N/A | 15% by wt | N/A |
| Triethanolamine | Thermo Fisher Scientific | 10% by wt | T407-500 |
| D-sucrose | Thermo Fisher Scientific | 50% by wt | S0389 |
| Urea | Sigma-Aldrich | 25% by wt | 15604 |

**Table 11.** Blocking solution composition.

| Reagents | Supplier/manufacturer | Working concentration | Catalogue number |
| --- | --- | --- | --- |
| Bovine Serum Albumin (BSA) | Wisent Bioproducts | 0.05% | 800-095-CG |
| Normal Donkey Serum (NDS) | Millipore | 5% | S30 |
| Phosphate Buffer Saline (PBS) | Wisent Bioproducts | 1X | 311-010-CL |
| Triton X-100 | Bioshop | 0.2% | TRX506 |

**Table 12.** List of consumables and equipment.

| Consumables and equipment | Supplier | Catalogue number |
| --- | --- | --- |
| 0.6 mL locking-lid microcentrifuge tubes | Thermo Fisher Scientific | 02-681-273 |
| Benchtop shaking incubator | Corning | LSE™ 6790 |
| Black bottom 96 well plate | Corning | 353219 |
| Nutating mixer | VWR | 82007-202 |
| Opera Phenix High-Content Screening System | PerkinElmer | N/A |
| Sterile cell strainer 40 µm | Thermo Fisher Scientific | 22363547 |
| Wide orifice low binding tips | Labcon | 1164-965-008-9 |

**Table 13.** Primary antibodies.

| Antibody | Host species | Working dilution | Reference |
| --- | --- | --- | --- |
| ChAT | Goat | 1:100 | Millipore; Cat. No. MAB144P |
| Hb9 (81.5C10) | Mouse | 1:50 | DSHB |
| Islet-1 (40.2D6) | Mouse | 1:50 | DSHB |
| Ki67 | Mouse | 1:200 | BD Biosciences; Cat. No. 556003 |
| Mbp | Rat | 1:200 | Novusbio; Cat. No. NB600-717 |
| Nestin | Mouse | 1:250 | Abcam; Cat. No. ab92391 |
| Neurofilament-H | Chicken | 1:1000 | Abcam; Cat. No. ab4680 |
| Nkx2.2 | Mouse | 1:100 | DSHB |
| Nkx6.1 | Mouse | 1:100 | DSHB |
| O4* | Mouse | 1:200 | R&D System; Cat. No. MAB1326 |
| Oct3/4 | Goat | 1:500 | Santa Cruz; Cat. No. sc-8628 |
| Olig2 | Rabbit | 1:100 | Millipore; Cat. No. AB 9610 |
| Pax6 | Mouse | 1:100 | DSHB |
| SMI-32 | Mouse | 1:100 | Biolegend; Cat. No. 801701 |
| Sox1 | Goat | 1:100 | R&D System; Cat. No. AF3369 |
| Uncx | Rabbit | 1:250 | Novusbio; Cat. No. NBP2-56480 |
| Vsx2 (Chx10) | Mouse | 1:250 | Santa-Cruz; Cat. No. sc-365519 |

\**Note: To obtain optimal results with O4 antibody, perform a live staining for 30 min at 37°C before formaldehyde fixation.*

**Table 14.** Secondary antibodies.

| Host species | Target species | Fluorophore | Working dilution | Reference |
| --- | --- | --- | --- | --- |
| Donkey | Goat IgG | DyLight 550 | 1:250 | Abcam; Cat. No. ab96932 |
| Donkey | Goat IgG | AlexaFluor 647 | 1:250 | Jackson ImmunoResearch; Cat. No. 703-605-155 |
| Donkey | Mouse IgG | DyLight t488 | 1:250 | Abcam; Cat. No. ab96875 |
| Donkey | Chicken IgG | AlexaFluor 647 | 1:250 | Jackson ImmunoResearch; Cat. No. 703-605-155 |
| Donkey | Rabbit IgG | DyLight 550 | 1:250 | Abcam; Cat. No. ab96892 |
| Donkey | Rabbit IgG | DyLight 488 | 1:250 | Abcam; Cat. No. ab96891 |
| Goat | Rat IgG | DyLight 488 | 1:250 | Abcam; Cat. No. ab96887 |

A maximum of six MN spheroids per 0.6 mL collection tube were transferred from the 96 U-bottom ultra-low attachment plates, fixed in 4% FA for 15-20 min at RT and washed three times with 1X PBS. To perform all the PBS washes of section **3.4**., the spheres were allowed to reach the bottom of the collection tube through gravity (~5 min) before removing the supernatant. For each wash, 400 μl of 1X PBS were used and the tube was left in the nutating mixer for 10 min. After fixation, the CUBIC protocol was performed by replacing 1X PBS by 200 μl of CUBIC R1. Incubation of the samples with CUBIC R1 was performed at 37°C with gentle shaking (~80 rpm) for 48-72 h. The removal of CUBIC R1 was achieved by performing three 1X PBS washes. MN spheroids were blocked in 200 μl of blocking solution overnight at 37°C and gentle shaking (~80 rpm). Blocking solution was removed by allowing the spheres to reach the bottom of the collection tube by gravity (~5 min). Primary antibodies were diluted in blocking solution and added to the MN spheroids for 24-72 h at 37°C and gentle shaking (~80 rpm). A final volume of 150 μl was used per collection. Primary antibodies were washed out by performing three 1X PBS washes. Secondary antibodies and Hoechst33342 were diluted in blocking solution and added to the MN spheroids for 24-72h at 37°C and gentle shaking (~80 rpm). A final volume of 150 μl was used per collection tube. Secondary antibodies and Hoechst33342 were washed out by performing three 1X PBS washes.

For imaging, we optimized a protocol to image MN spheroids at large scale using black 96 well plates. Each spheroid was transferred into the center of a well using wide orifice low binding tips. Excess 1X PBS was removed and 100 μl of CUBIC R2 was added per well as mounting medium. Images of the immunostained MN spheroids were acquired with the Opera Phenix High-Content Screening System using the PreScan function to find the spheres within the focal plane at 5X and then perform the imaging at 20X. System 5x/0.16 and 20X/1.0 objectives. ‘Image size 512 × 512, voxel size 2.9 × 2.9 × 5 µm. The data was extracted to be organized and analyzed by an in-house script developed in MATLAB. Alternatively, MN spheroids can be mounted over a glass slide to be imaged with a conventional confocal microscope. For this, a hydrophobic pen is used to draw a circle in the middle of a glass slide and a MN spheroid is transferred in the center using wide orifice low binding tips. Excess 1X PBS was removed and a drop of CUBIC R2 (~30 μl) is added to the sphere followed by the placing of a glass coverslip on top.

**Note:** CUBIC R1 and R2 solutions must be filtered using a cell strainer to remove any solutes that did not incorporate into the solution. This will reduce undesired detritus at the imaging step. After the addition of CUBIC R1, the spheroids become transparent, and they are almost imperceptible to the eye. After the first 1X PBS wash, the spheres recover their white color and it is possible to see them again. For the mounting process inside black 96 well plates, it is essential to remove almost all the 1X PBS surrounding the spheroids before adding the CUBIC R2. This will prevent the spheroids from floating at different heights inside the well, thus making their tracing difficult at the microscope.

### 3.5 Microelectrode array (MEA) recordings of MN spheroids

Each batch of MNPCs was used at passage 2 to generate one 96 U-bottom ultra-low attachment plates (60 spheroids). The material, reagents, and equipment to perform MN spheroid MEA recordings are listed in **Table 15**. The consumables and equipment are listed in **Table 16**. The composition of the artificial cerebrospinal fluid (aCSF) solution needed for MEA recordings is detailed in **Table 17**.

**Table 15.** List of material and reagents.

| Reagents | Supplier/manufacturer | Working concentration | Catalogue number |
| --- | --- | --- | --- |
| Laminin | Invitrogen | 5 µg/mL | L2020 |
| Poly-L-ornithine (PLO) | Sigma-Aldrich | 10 µg/mL | P3655 |
| Tetrodotoxin (TTX) | Sigma-Aldrich | 1 mM | T8024 |

**Table 16.** List of consumables and equipment.

| Consumables and equipment | Supplier | Catalogue number |
| --- | --- | --- |
| 0.2 µm filters | Thermo Fisher Scientific | 09-719C |
| 30 mL Syringe | VWR | 76290-386 |
| 50 mL conical tube | Progene | 71-5000-B |
| Axion Maestro Edge | Axion Biosystems | N/A |
| Cytoview 24-well MEA plates | Axion Biosystems | M384-Tmea-24w |
| Wide orifice low binding tips | Labcon | 1164-965-008-9 |

**Table 17.** Composition of aCSF.

| Reagents** | Supplier/manufacturer | Working concentration | Catalogue number |
| --- | --- | --- | --- |
| CaCl<sub>2</sub> | Thermo Fisher Scientific | 1.6 mM | AC423525000 |
| D-glucose* | Sigma-Aldrich | 5.5 mM | G8270 |
| KCl | Bioshop | 4 mM | POC888.5 |
| KH<sub>2</sub>PO<sub>4</sub> | Thermo Fisher Scientific | 1.18 mM | AC424200250 |
| MgSO<sub>4</sub> | Thermo Fisher Scientific | 1.17 mM | AC423905000 |
| NaCl | BioShop | 119 mM | SOD004.5 |
| NaHCO<sub>3</sub>* | Thermo Fisher Scientific | 24 mM | AC424270010 |

\* These reagents are not added to the 10X stock solution to prevent the growth of microorganisms. They are added to the 1X solution before every recording experiment.
\*\* Add 2g of NaCl to 10X solution stock for adjusting osmolality 305mOsm.

Cytoview 24-well MEA plates with 16 electrodes per well were treated with PLO (10 μg/ml) for 24 h, washed three times with 1X PBS and coated with laminin (5 μg/ml) for 24 h. MN spheroids were generated and cultured for 7 days into the 96 U-bottom ultra-low attachment well plates before being transferred to each well of the Cytoview 24-well MEA plate using wide orifice low binding tips. Six spheroids were deposited in the center of each well in no more than 20 μl of medium to prevent the spheroids from spreading to the edges of the well. The plate was returned to the incubator for 20-30 min to allow the spheroids attach to the electrodes before addition of 500 μl of fresh MN induction and maturation medium.

Before each recording, 50 mL of 1X aCSF solution was prepared from a 10X stock solution, adding D-glucose and NaHCO_3_ at the working concentrations. The tube with the 1X aCSF solution was left with the lid loose inside the incubator for 1 h to equilibrate the solution with the same gaseous conditions found inside the incubator. Before the 1X aCSF solution was added to the cells, it was filtered to avoid any possible contamination. After removing the MN induction and maturation medium and adding 500 μl of the 1X aCSF solution to each well, the MEA plate was returned to the incubator at 37°C/5% CO_2_ for 30-45 min.

MEA recordings were performed on day 7, 14, 21 and 28 after plating the MN spheroids. Data was collected for 5min using the Axis Navigator software provided by Axion Biosystems. A band-pass filter of 3 kHz (low-pass) to 200 Hz (high-pass) was applied. For the analysis, a “spike” was defined as a short extracellular electrical event with a peak voltage six times or greater than the standard deviation of the estimated “noise” signal. A “burst” was defined as ≤5 spikes with no more than 100ms separating each spike. Network bursts were not measured as the presence of spheres within the plate was random with very few spheres centered within an electrode for the entire recording period of 28 days. The MEA plate was placed into the Axion Maestro Edge with temperature and CO_2_ concentration set to 37°C and 5% respectively. The plate was allowed to equilibrate for 5 min inside the instrument prior to recording. After the recording, the MEA plate was removed from the instrument and the aCSF was replaced with MN induction and maturation medium to keep the cells in culture for the following recording time points. At day 28, after the basal recording of 5 min in aCSF was performed, the plate was removed from the instrument and dosed with vehicle (H_2_O) or 1mM tetrodotoxin. The plate was returned to the instrument and allowed to equilibrate for 5 min before performing a second recording of 5 min. This was the endpoint of the MEA recordings.

## 4 Results

We generated MN spheroids from two different healthy control iPSC lines (AIW002-02 and 3450), adapting a previously described protocol used to generate MNs as a 2D monolayer. For our experiments, 80% confluent iPSC cultures were differentiated into MNPCs. At the MNPC stage, we confirmed the expression of the neural precursor markers Sox1, Nestin and Pax6 (43, 44) in addition to Ki67, an endogenous marker of active cell cycle, thus confirming the cells proliferation capacity (45) (**Supplementary Figure 1**). MNPCs also co-expressed the markers Nkx6.1 and Olig2, confirming their identity as MNPCs (46) (**Supplementary Figure 1**). From these MNPCs, 5,000 cells per well were used to generate iPSC-derived MN spheroids into 96-well U-bottom ultra-low attachment plates (**Fig 1B**). Diameter (3450, 14 days= 262.9 μm ± 1.976, 28 days=278.3 μm ± 2.255; AIW002-02, 14 days= 305.4 μm ± 2.225, 28 days=326.4 μm ± 1.752), area (3450, 14 days= 5.92×10^4^ μm^2^ ± 804.4, 28 days=6.61×10^4^ μm^2^ ± 1065; AIW002-02, 14 days= 8.13 ×10^4^ μm^2^ ± 1090, 28 days=9.40 ×10^4^ μm^2^ ± 1244) and volume (3450, 14 days= 9.73 ×10^6^ μm^3^ ± 214272, 28 days=1.15×10^7^ μm^3^ ± 274041; AIW002-02, 14 days= 1.52 ×10^7^ μm^3^ ± 346330, 28 days=1.84 ×10^7^ μm^3^ ± 303753) measurements of the MN spheroids were obtained using Cell Profiler Analyst. We observed that the spheroids from both control lines grew in size from 14 to 28 days as confirmed by increases in the measurements of their overall diameter (3450= p<0.0001; AIW002-02=p<0.0001), area (3450= p<0.0001; AIW002-02=p<0.0001) and volume (3450= p<0.0001; AIW002-02=p<0.0001) (**Fig 2A/B**). Importantly, since our protocol allows us to expand the MNPCs for up to 5 passages (13), we generated MN spheroids from MNPCs between passages 2 to 4 and we observed that the spheroids were successfully generated and kept their capacity to grow over time regardless of the passage number. In terms of morphology, the MN spheroids display a circular shape, however, an analysis of circularity (**Fig 2C**) showed that the spheroids are not perfectly round. Taken together, we succeeded in the establishment of culture conditions to generate 3D iPSC-derived MN spheroids in a reproducible manner.

Next, we assessed the expression of MN markers in the MN spheroids derived from the two control lines (AIW002-02 and 3450) at the transcript and protein level by qPCR (**Fig 3**) and immunofluorescent staining (**Fig 4**) respectively. For both 3450 (**Fig 3A**) and AIW002-02 (**Fig 3B**) lines, the expression of different genes at the transcript level was assessed at the iPSC (NANOG and OCT4), MNPC (OLIG2 and PAX6) and MN spheroid (HB9, ISL1, CHAT and MAP2) stage. As expected, the pluripotency markers NANOG and OCT4 were found to be downregulated at the MNPC and MN spheroid (14 and 18 days) stage in both cell lines. The MNPC markers PAX6 and OLIG2 were found upregulated at the MNPC stage, while being downregulated at the iPSC and MN spheroid (14 and 28) stage in both cell lines. Finally, the expression level of the neural marker MAP2, as well as the expression levels of the MN markers HB9, ISL1 and CHAT were found to be upregulated at the MN spheroid stage while being almost absent at the iPSC and MNPC stages in both cell lines. Additionally, we assessed the expression levels of interneuron (VSX2 and SIM1), oligodendrocyte (MBP) and astrocyte (GFAP) markers to address the presence of other cell types within the MN spheroids. For both cell lines, we confirmed the upregulation of MN markers at the transcript level, indicating a successful differentiation towards MN identity. GFAP expression remained undetected at the different differentiation stages indicating the absence of astrocytes. However, we observed the expression of interneuron and oligodendrocyte markers suggesting the presence of these cell types within the MN spheroids.

**Figure 3.**
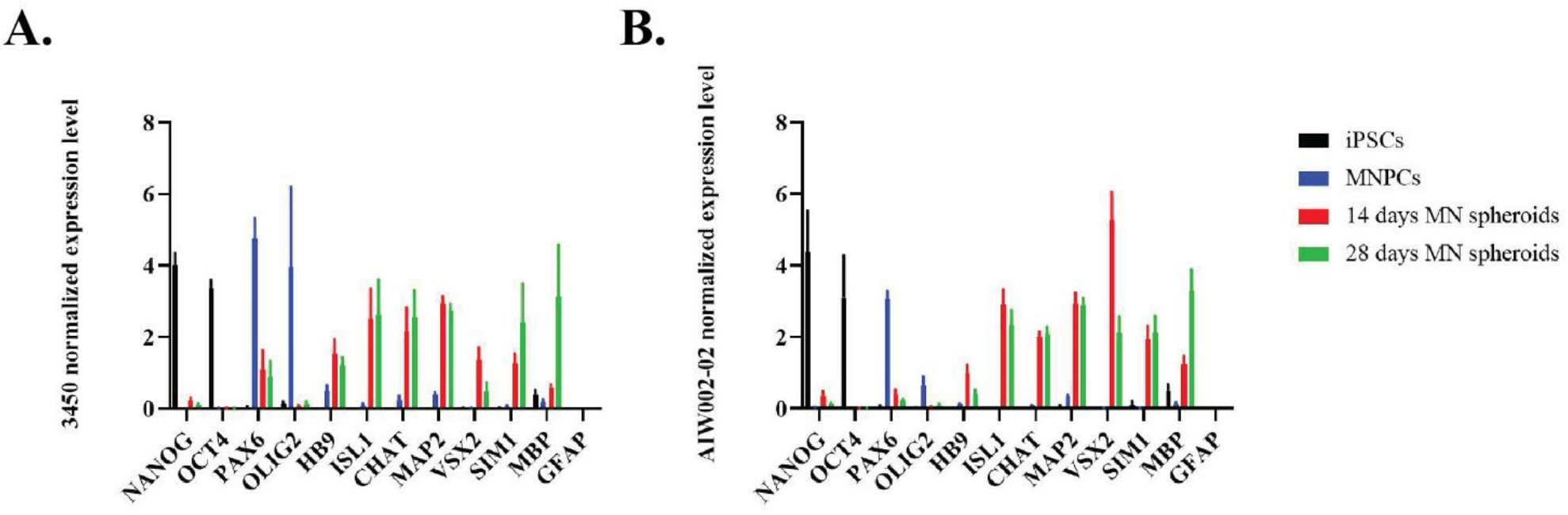
Transcript expression profile of iPSC-derived MN spheroids. Normalized expression levels of NANOG, OCT4, PAX6, OLIG2, HB9, ISL1, CHAT MAP2, VSX2, SIM1, MBP and GFAP in iPSCs, MNPCs, and iPSC-derived MN spheroids differentiated for 14 and 28 days from **A)** 3450 and **B)** AIW002-02 lines. Data normalized to Actβ-GAPDH expression. Bar graphs show the mean ± SEM; three MN spheroid batches per time point (14 and 28 days). Each MN spheroid batch was obtained from iPSC-derived MNPC batches at passage 3 generated through independent differentiation processes.

**Figure 4.**
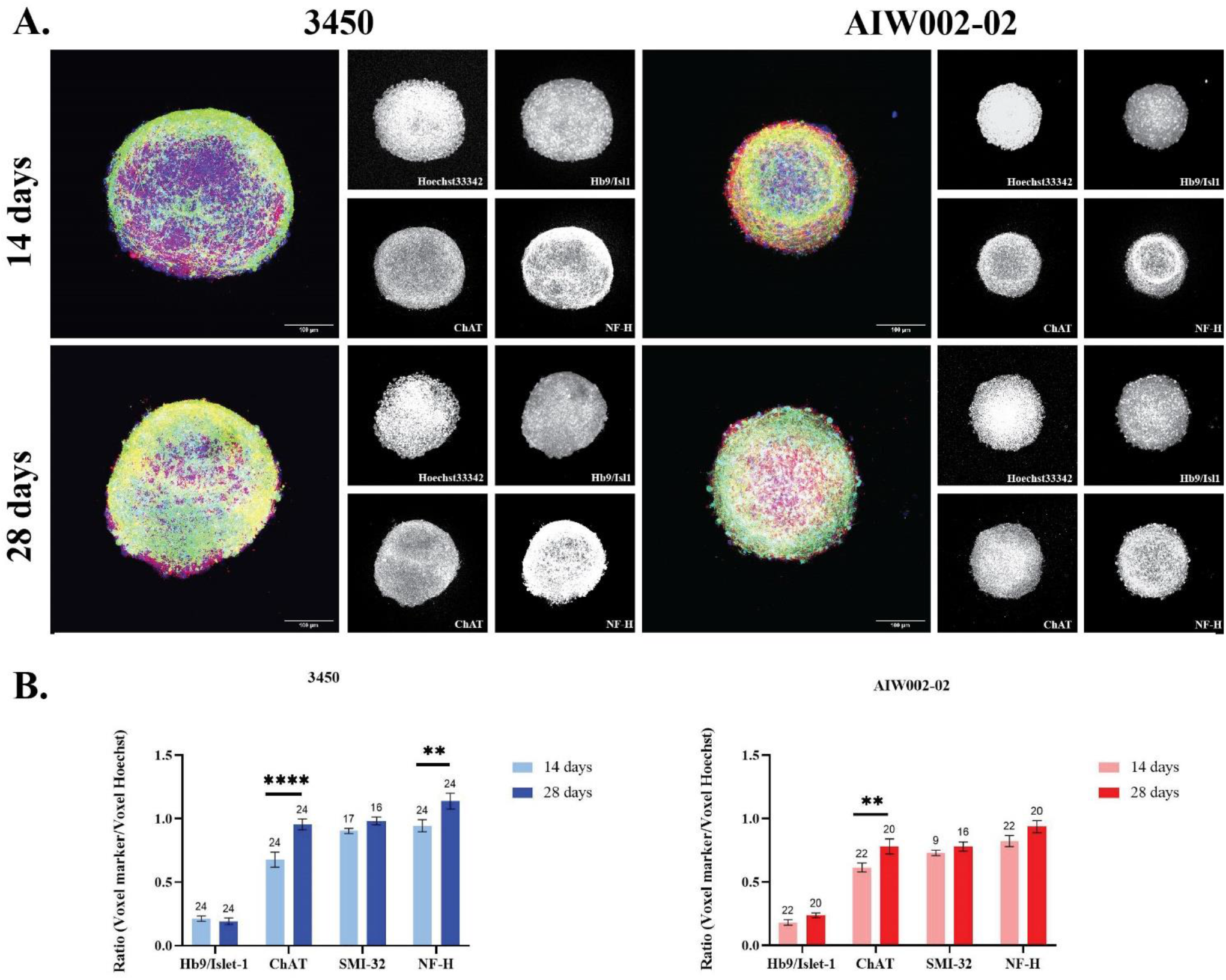
Image profiling of iPSC-derived MN spheroids. **A)** Characterization of MN spheroids by immunostaining showed that MN spheroids generated from the 3450 and AIW002-02 iPSC lines express the specific MN markers Hb9, Islet-1, ChAT and SMI-32 (not shown) as well as the neuronal marker NF-H. Merge shows overlay of Hb9/Islet-1 (blue), ChAT (red), and NF-H (green). **B)** The presence of each MN marker was quantified using an in-house MATLAB pipeline. Graph bars show the mean ± SEM; for each cell line, each batch of three batches of iPSC-derived MNPCs generated through independent differentiation processes were used to generate two MN spheroid batches. A minimum of 9 MN spheroids were required for quantification, and we ensured that at least 3 spheroids per MNPC batch where stained for each marker per cell line at each time point. The significance between the two different time points (14 days and 28 days) for each MN marker was determined by a unpaired *t*-student **p<0.01; ****p<0.0001.

To determine the presence of the different markers at the protein level we analyzed the images of immunostained MN spheroids at 14 and 28 days with our in-house MATLAB script. We confirmed the expression of the pan MN markers Hb9/Isl1 after 14 and 28 days of differentiation with no statistical difference observed at the different time points for either cell line (**Fig 4B**). The expression of ChAT, a marker associated to MN maturation, was also assessed, showing an increase of its expression at 28 days compared to 14 days for both cells lines (3450= p<0.0001; AIW002-02=p<0.01) (**Fig 4B**). The expression of neurofilament heavy (NF-H), a marker indicating neuronal identity, was also increased at 28 days in both cell lines, however, it only reached a statistical significance in one of the cells lines (3450= p<0.01) (**Fig 4B**). Finally, we assessed the expression of SMI-32, an antibody that recognizes the non-phosphorylated form of the neurofilament medium (NF-M) and NF-H subunits, which is known to be highly enriched in spinal motor neurons. The expression of SMI-32 was confirmed at the two time points for both cell lines. We observe a tendency of SMI-32 to increase its expression at 28 days; however, it did not reach the statistical significance for either of the cell lines (**Fig 4B**). MN spheroids were also stained for pluripotency (Oct3/4) and MNPC (Pax6 and Olig2) markers to confirm their downregulation (**Supplementary Figure 3**). The expression of Hb9, Isl1, ChAT and NF-H, confirmed that at the protein level, the MN spheroids are composed primarily of neurons with a MN identity. Nevertheless, positive antibody staining for interneuron progenitor (NKX2.2), interneuron (Vsx2 and Uncx) and oligodendrocyte (O4 and MBP) markers (**Supplementary Figure 3**), confirms the presence of other cell types in the MN spheroids, as suggested by the qPCR analysis (**Fig 4C**).

Finally, we used the MEA system to assess the electrical activity of the MN spheroids. Only the spheroids placed at the center of the electrode were considered for quantification, meaning that each spheroid was in contact with a single electrode within the well (**Fig 5A**). Action potentials (“spikes”) and groups of action potentials (“bursts”) were detected for the MN spheroids derived from the two control cell lines (3450 and AIW002-02) at different time points (7, 14, 21 and 28 days). The mean firing rate (Hz), a ratio of the total number of spikes recorded over the duration of the recording, indicates that MN spheroids remain electrically active over time as previously described for 2D monolayer iPSC-derived MNs MEA recordings (47) (**Fig 5B**). Additionally, MN spheroids display burst firing over time (**Fig 5C**). Finally, treatment with TTX, a sodium channel blocker that inhibits the firing of action potential in neurons, confirmed the electrical activity of the MN spheroids ruling out any potential artifacts coming from the system. Activity was not altered when a well was treated with vehicle (**Fig 5D**).

**Figure 5.**
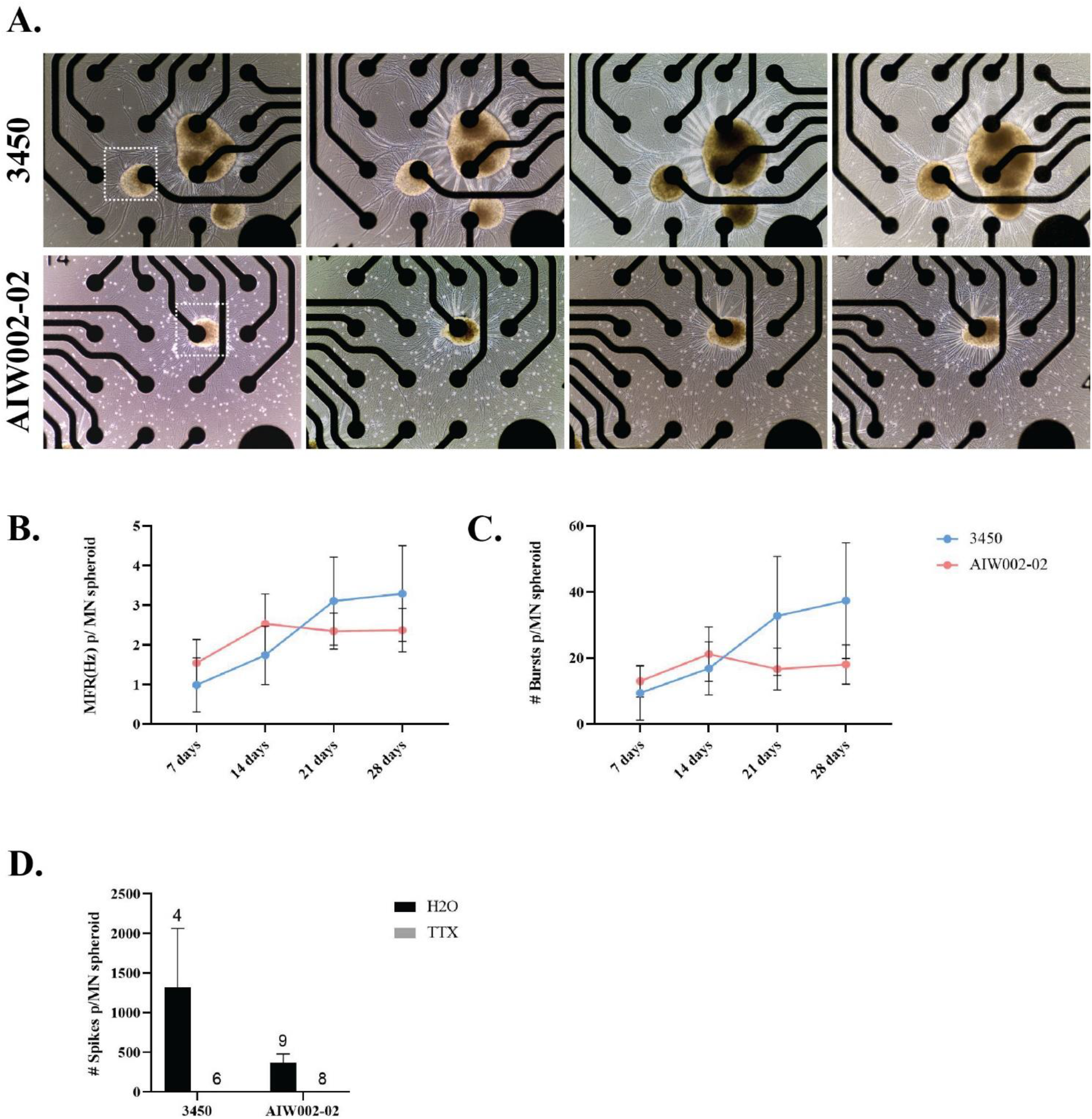
Spontaneous firing activity of iPSC-derived MN spheroids at different time points. **A)** iPSC-derived MN spheroids were plated on wells of Cytoview 24-well MEA plates and recorded every 7 days until 28 days. Only MN spheroids placed at the center of an electrode (squares) through the entire recording time were considered for quantification. **B)** Mean Firing Rate (MFR) and **C)** number# of bursts were plotted per individual MN spheroid. The connected scatter plots show the mean ± SEM; for each cell line, each batch of three batches of iPSC-derived MNPCs generated through independent differentiation processes were used to generate one MN spheroid batch. A minimum of nine MN spheroids were required for quantification to ensure that at least 3 spheroids per MNPC batch were quantified. **D)** Tetrodotoxin (TTX) treatment blocked all firing in MN spheroids ruling out system artifacts and confirming neural activity for both cell lines. H_2_O was used as vehicle. Graph bars show mean ±SEM.

All graphs presented in section **4** were generated using GraphPad Prism version 9.1.1 for Windows, GraphPad Software, San Diego, California USA, www.graphpad.com.

## 5 Discussion

Taken together, the results presented here demonstrate that we were able to develop a workflow to generate and characterize MN spheroids from iPSCs by modifying a protocol originally optimized to generate a 2D MN monolayer culture. Culturing the cells in a 3D context did not interfere with the expression of MN identity markers as shown by qPCR and immunofluorescent stainings. However, we found the presence of interneuron and oligodendrocyte markers within the MN spheroids which is consistent with observations from our single-cell RNA sequencing (48) when iPSC-derived MNs were cultured as a 2D monolayer following the original protocol (13). Therefore, this needs to be taken into consideration when assays are performed in these 3D models to avoid misinterpretations of results.

Even in small 3D samples like spheroids, imaging is challenging due to the intrinsic opacity of biological tissues. Tissue clearing techniques are useful to reduce the water content of a sample and to remove lipids harbored within its cell membranes to 1) facilitate antibody penetrance and immunostaining, and 2) reduce microscope light scattering to improve 3D image acquisition. Here, we were able to adapt a previously described water-based clearing protocol, CUBIC (42, 49, 50), to clear MN spheroids. The performance of this technique improved the immunofluorescent staining and subsequent image acquisition of our samples. When the CUBIC protocol was not performed, antibody penetration was limited to the periphery of the spheroid making its center perceived as a necrotic core.

MEA recordings of the MN spheroids confirmed the presence of neuronal activity within these 3D structures as demonstrated by their ability to fire action potentials as isolated spikes or in bursts. Burst patterns measurements obtained by performing MEA recordings have been used to assess electrical changes in 2D monolayer cortical cultures (51), however, to our knowledge there are not equivalent studies using MN cultures. Thus, this assay opens the possibility of using a MEA approach to analyze differences in burst patterns when MN spheroids are derived from control vs patient iPSC lines. Nevertheless, considering the presence of other cell types within the MN spheroids, it would be important to identify the different types of synapses present in these 3D structures by applying ion channel-specific antagonists. From the technical perspective, placing the MN spheroids at the center of a single electrode is challenging. Therefore, for future functional characterization, we acknowledge that a better approach would be the use of high-density MEAs (HD-MEAs) which have been recently used to assess the electrical features of other 3D structures (52, 53). HD-MEAs are composed of thousands of electrodes with minimal space between them. Therefore, HD-MEAs systems not only provide an increase of spatiotemporal resolution but they would also facilitate the seeding process of 3D structures.

Even though there has been an increase in studies making use of MN spheroids, this is the first time a broader characterization has been made for such 3D structures with tools specifically developed for large-scale studies. This is relevant because iPSC-derived spheroids/neurospheres have proven to be easy to generate 3D models that can be used for screening purposes (54), an important step towards drug discovery for MNDs. Moreover, this model can be used as a cellular jigsaw, in which other iPSC-derived cell types can be added to form more advanced co-culture spheroids, opening the possibility to study non cell autonomous manifestation of the diseases.

## 7 Author Contributions

MJC-M: conception and design, data analysis, compiling and summarizing articles, figures elaboration, manuscript writing, final approval of the manuscript. MC: design, final approval of the manuscript. AKF-F: experimental data analysis, MNPC generation and design of the Cell Profiler pipeline. DC-V: data analysis. WER: design of the Cell Profiler pipeline. TMD: financial support, design, correction of style, manuscript writing, final approval of the manuscript.

## 9 Acknowledgments

We acknowledge Dr. Veronica García-Vázquez and Dr. Maria Lacalle-Aurioles for the design and coding of the in-house developed MATLAB script. We acknowledge Ghislaine Deyab for her guidance with MEA data analysis. We thank Sarah Lépine, Dr. Lenore K. Beitel and Dr. Mark Aurousseau for editing and proofreading of the manuscript.

## References

1. Sanes J.R. & Lichtman J.W. Development of the vertebrate neuromuscular junction. Annu Rev Neurosci. 1999;22:389–442.

2. Tiryaki E. & Horak H.A. ALS and other motor neuron diseases. Continuum (Minneap Minn). 2014;20(5 Peripheral Nervous System Disorders):1185–207.

3. Sances S., Bruijn L.I., Chandran S., Eggan K., Ho R., Klim J.R., Livesey M.R., Lowry E., Macklis J.D., Rushton D., Sadegh C., Sareen D., Wichterle H., Zhang S.C. & Svendsen C.N. Modeling ALS with motor neurons derived from human induced pluripotent stem cells. Nat Neurosci. 2016;19(4):542–53.

4. Takahashi K. & Yamanaka S. Induction of pluripotent stem cells from mouse embryonic and adult fibroblast cultures by defined factors. Cell. 2006;126(4):663–76.

5. Takahashi K., Tanabe K., Ohnuki M., Narita M., Ichisaka T., Tomoda K. & Yamanaka S. Induction of pluripotent stem cells from adult human fibroblasts by defined factors. Cell. 2007;131(5):861–72.

6. Fujimori K., Ishikawa M., Otomo A., Atsuta N., Nakamura R., Akiyama T., Hadano S., Aoki M., Saya H., Sobue G. & Okano H. Modeling sporadic ALS in iPSC-derived motor neurons identifies a potential therapeutic agent. Nat Med. 2018;24(10):1579–89.

7. Ratti A., Gumina V., Lenzi P., Bossolasco P., Fulceri F., Volpe C., Bardelli D., Pregnolato F., Maraschi A., Fornai F., Silani V. & Colombrita C. Chronic stress induces formation of stress granules and pathological TDP-43 aggregates in human ALS fibroblasts and iPSC-motoneurons. Neurobiol Dis. 2020;145:105051.

8. Kim B.W., Ryu J., Jeong Y.E., Kim J. & Martin L.J. Human motor neurons with SOD1-G93A mutation generated from CRISPR/Cas9 gene-edited iPSCs develop pathological features of amyotrophic lateral sclerosis. Front Cell Neurosci. 2020;14:604171.

9. Deneault E., Chaineau M., Nicouleau M., Castellanos Montiel M.J., Franco Flores A.K., Haghi G., Chen C.X., Abdian N., Shlaifer I., Beitel L.K. & Durcan T.M. A streamlined CRISPR workflow to introduce mutations and generate isogenic iPSCs for modeling amyotrophic lateral sclerosis. Methods. 2021.

10. Selvaraj B.T., Livesey M.R., Zhao C., Gregory J.M., James O.T., Cleary E.M., Chouhan A.K., Gane A.B., Perkins E.M., Dando O., Lillico S.G., Lee Y.B., Nishimura A.L., Poreci U., Thankamony S., Pray M., Vasistha N.A., Magnani D., Borooah S., Burr K., Story D., McCampbell A., Shaw C.E., Kind P.C., Aitman T.J., Whitelaw C.B.A., Wilmut I., Smith C., Miles G.B., Hardingham G.E., Wyllie D.J.A. & Chandran S. C9ORF72 repeat expansion causes vulnerability of motor neurons to Ca(2+)-permeable AMPA receptor-mediated excitotoxicity. Nat Commun. 2018;9(1):347.

11. Guo W., Naujock M., Fumagalli L., Vandoorne T., Baatsen P., Boon R., Ordovás L., Patel A., Welters M., Vanwelden T., Geens N., Tricot T., Benoy V., Steyaert J., Lefebvre-Omar C., Boesmans W., Jarpe M., Sterneckert J., Wegner F., Petri S., Bohl D., Vanden Berghe P., Robberecht W., Van Damme P., Verfaillie C. & Van Den Bosch L. HDAC6 inhibition reverses axonal transport defects in motor neurons derived from FUS-ALS patients. Nat Commun. 2017;8(1):861.

12. Boulting G.L., Kiskinis E., Croft G.F., Amoroso M.W., Oakley D.H., Wainger B.J., Williams D.J., Kahler D.J., Yamaki M., Davidow L., Rodolfa C.T., Dimos J.T., Mikkilineni S., MacDermott A.B., Woolf C.J., Henderson C.E., Wichterle H. & Eggan K. A functionally characterized test set of human induced pluripotent stem cells. Nat Biotechnol. 2011;29(3):279–86.

13. Du Z.W., Chen H., Liu H., Lu J., Qian K., Huang C.L., Zhong X., Fan F. & Zhang S.C. Generation and expansion of highly pure motor neuron progenitors from human pluripotent stem cells. Nat Commun. 2015;6:6626.

14. Qu Q., Li D., Louis K.R., Li X., Yang H., Sun Q., Crandall S.R., Tsang S., Zhou J., Cox C.L., Cheng J. & Wang F. High-efficiency motor neuron differentiation from human pluripotent stem cells and the function of Islet-1. Nat Commun. 2014;5:3449.

15. Amoroso M.W., Croft G.F., Williams D.J., O’Keeffe S., Carrasco M.A., Davis A.R., Roybon L., Oakley D.H., Maniatis T., Henderson C.E. & Wichterle H. Accelerated high-yield generation of limb-innervating motor neurons from human stem cells. J Neurosci. 2013;33(2):574–86.

16. Dimos J.T., Rodolfa K.T., Niakan K.K., Weisenthal L.M., Mitsumoto H., Chung W., Croft G.F., Saphier G., Leibel R., Goland R., Wichterle H., Henderson C.E. & Eggan K. Induced pluripotent stem cells generated from patients with ALS can be differentiated into motor neurons. Science. 2008;321(5893):1218–21.

17. Maury Y., Côme J., Piskorowski R.A., Salah-Mohellibi N., Chevaleyre V., Peschanski M., Martinat C. & Nedelec S. Combinatorial analysis of developmental cues efficiently converts human pluripotent stem cells into multiple neuronal subtypes. Nat Biotechnol. 2015;33(1):89–96.

18. Karpe Y., Chen Z. & Li X.J. Stem cell models and gene targeting for human motor neuron diseases. Pharmaceuticals (Basel). 2021;14(6).

19. Guo W., Fumagalli L., Prior R. & Van Den Bosch L. Current advances and limitations in modeling ALS/FTD in a dish using induced pluripotent stem cells. Front Neurosci. 2017;11:671.

20. Afshar Bakooshli M., Lippmann E.S., Mulcahy B., Iyer N., Nguyen C.T., Tung K., Stewart B.A., van den Dorpel H., Fuehrmann T., Shoichet M., Bigot A., Pegoraro E., Ahn H., Ginsberg H., Zhen M., Ashton R.S. & Gilbert P.M. A 3D culture model of innervated human skeletal muscle enables studies of the adult neuromuscular junction. Elife. 2019;8.

21. de Leeuw S.M., Davaz S., Wanner D., Milleret V., Ehrbar M., Gietl A. & Tackenberg C. Increased maturation of iPSC-derived neurons in a hydrogel-based 3D culture. J Neurosci Methods. 2021;360:109254.

22. Brännvall K., Bergman K., Wallenquist U., Svahn S., Bowden T., Hilborn J. & Forsberg-Nilsson K. Enhanced neuronal differentiation in a three-dimensional collagen-hyaluronan matrix. J Neurosci Res. 2007;85(10):2138–46.

23. Smith I., Haag M., Ugbode C., Tams D., Rattray M., Przyborski S., Bithell A. & Whalley B.J. Neuronal-glial populations form functional networks in a biocompatible 3D scaffold. Neurosci Lett. 2015;609:198–202.

24. Vagaska B., Gillham O. & Ferretti P. Modelling human CNS injury with human neural stem cells in 2- and 3-Dimensional cultures. Sci Rep. 2020;10(1):6785.

25. Thiry L., Clément J.P., Haag R., Kennedy T.E. & Stifani S. Optimization of long-term human iPSC-derived spinal motor neuron culture using a dendritic polyglycerol amine-based substrate. ASN Neuro. 2022;14:17590914211073381.

26. Taga A., Dastgheyb R., Habela C., Joseph J., Richard J.P., Gross S.K., Lauria G., Lee G., Haughey N. & Maragakis N.J. Role of human-induced pluripotent stem cell-derived spinal cord astrocytes in the functional maturation of motor neurons in a multielectrode array system. Stem Cells Transl Med. 2019;8(12):1272–85.

27. Toli D., Buttigieg D., Blanchard S., Lemonnier T., Lamotte d’Incamps B., Bellouze S., Baillat G., Bohl D. & Haase G. Modeling amyotrophic lateral sclerosis in pure human iPSc-derived motor neurons isolated by a novel FACS double selection technique. Neurobiol Dis. 2015;82:269–80.

28. Faustino Martins J.M., Fischer C., Urzi A., Vidal R., Kunz S., Ruffault P.L., Kabuss L., Hube I., Gazzerro E., Birchmeier C., Spuler S., Sauer S. & Gouti M. Self-organizing 3D human trunk neuromuscular organoids. Cell Stem Cell. 2020;26(2):172-86.e6.

29. Pereira J.D., DuBreuil D.M., Devlin A.C., Held A., Sapir Y., Berezovski E., Hawrot J., Dorfman K., Chander V. & Wainger B.J. Human sensorimotor organoids derived from healthy and amyotrophic lateral sclerosis stem cells form neuromuscular junctions. Nat Commun. 2021;12(1):4744.

30. Hor J.H. & Ng S.Y. Generating ventral spinal organoids from human induced pluripotent stem cells. Methods Cell Biol. 2020;159:257–77.

31. Andersen J., Revah O., Miura Y., Thom N., Amin N.D., Kelley K.W., Singh M., Chen X., Thete M.V., Walczak E.M., Vogel H., Fan H.C. & Paşca S.P. Generation of functional human 3D cortico-motor assembloids. Cell. 2020;183(7):1913-29.e26.

32. Osaki T., Uzel S.G.M. & Kamm R.D. On-chip 3D neuromuscular model for drug screening and precision medicine in neuromuscular disease. Nat Protoc. 2020;15(2):421–49.

33. Machado C.B., Pluchon P., Harley P., Rigby M., Gonzalez Sabater V., Stevenson D.C., Hynes S., Lowe A., Burrone J., Viasnoff V. & Lieberam I. In vitro modelling of nerve-muscle connectivity in a compartmentalised tissue culture device. Adv Biosyst. 2019;3(7).

34. Kawada J., Kaneda S., Kirihara T., Maroof A., Levi T., Eggan K., Fujii T. & Ikeuchi Y. Generation of a motor nerve organoid with human stem cell-derived neurons. Stem Cell Reports. 2017;9(5):1441–9.

35. Yamamoto K., Yamaoka N., Imaizumi Y., Nagashima T., Furutani T., Ito T., Okada Y., Honda H. & Shimizu K. Development of a human neuromuscular tissue-on-a-chip model on a 24-well-plate-format compartmentalized microfluidic device. Lab Chip. 2021;21(10):1897–907.

36. Osaki T., Uzel S.G.M. & Kamm R.D. Microphysiological 3D model of amyotrophic lateral sclerosis (ALS) from human iPS-derived muscle cells and optogenetic motor neurons. Sci Adv. 2018;4(10):eaat5847.

37. Chen X., Rocha, Cecilia, Rao, Trisha, & Durcan, Thomas M.. Motor neuron induction and differentiation (V2.0). Zenodo. 2019.

38. Chen C., Abdian N., Maussion G., Thomas R.A., Demirova I., Cai E., Tabatabaei M., K. B.L., Karamchandani J., A. F.E. & M. D.T. Standardized quality control workflow to evaluate the reproducibility and differentiation potential of human iPSCs into neurons. bioRxiv. 2021.

39. Yoshida T. & Delafontaine P. Mechanisms of IGF-1-mediated regulation of skeletal muscle hypertrophy and atrophy. Cells. 2020;9(9).

40. Oyesiku N.M. & Wigston D.J. Ciliary neurotrophic factor stimulates neurite outgrowth from spinal cord neurons. J Comp Neurol. 1996;364(1):68–77.

41. Jones T.R., Kang I.H., Wheeler D.B., Lindquist R.A., Papallo A., Sabatini D.M., Golland P. & Carpenter A.E. CellProfiler Analyst: data exploration and analysis software for complex image-based screens. BMC Bioinformatics. 2008;9:482.

42. Gómez-Gaviro M.V., Balaban E., Bocancea D., Lorrio M.T., Pompeiano M., Desco M., Ripoll J. & Vaquero J.J. Optimized CUBIC protocol for three-dimensional imaging of chicken embryos at single-cell resolution. Development. 2017;144(11):2092–7.

43. Lendahl U., Zimmerman L.B. & McKay R.D. CNS stem cells express a new class of intermediate filament protein. Cell. 1990;60(4):585–95.

44. Neely M.D., Litt M.J., Tidball A.M., Li G.G., Aboud A.A., Hopkins C.R., Chamberlin R., Hong C.C., Ess K.C. & Bowman A.B. DMH1, a highly selective small molecule BMP inhibitor promotes neurogenesis of hiPSCs: comparison of PAX6 and SOX1 expression during neural induction. ACS Chem Neurosci. 2012;3(6):482–91.

45. Gerdes J., Lemke H., Baisch H., Wacker H.H., Schwab U. & Stein H. Cell cycle analysis of a cell proliferation-associated human nuclear antigen defined by the monoclonal antibody Ki-67. J Immunol. 1984;133(4):1710–5.

46. Ogura T., Sakaguchi H., Miyamoto S. & Takahashi J. Three-dimensional induction of dorsal, intermediate and ventral spinal cord tissues from human pluripotent stem cells. Development. 2018;145(16).

47. Ronchi S., Buccino A.P., Prack G., Kumar S.S., Schröter M., Fiscella M. & Hierlemann A. Electrophysiological phenotype characterization of human iPSC-derived neuronal cell lines by means of high-density microelectrode arrays. Adv Biol (Weinh). 2021;5(3):e2000223.

48. Thiry L., Hamel R., Pluchino S., Durcan T. & Stifani S. Characterization of human iPSC-derived spinal motor neurons by single-cell RNA sequencing. Neuroscience. 2020;450:57–70.

49. Susaki E.A., Tainaka K., Perrin D., Kishino F., Tawara T., Watanabe T.M., Yokoyama C., Onoe H., Eguchi M., Yamaguchi S., Abe T., Kiyonari H., Shimizu Y., Miyawaki A., Yokota H. & Ueda H.R. Whole-brain imaging with single-cell resolution using chemical cocktails and computational analysis. Cell. 2014;157(3):726–39.

50. Susaki E.A., Tainaka K., Perrin D., Yukinaga H., Kuno A. & Ueda H.R. Advanced CUBIC protocols for whole-brain and whole-body clearing and imaging. Nat Protoc. 2015;10(11):1709–27.

51. Wagenaar D.A., Pine J. & Potter S.M. An extremely rich repertoire of bursting patterns during the development of cortical cultures. BMC Neurosci. 2006;7:11.

52. Sharf T., van der Molen T., Glasauer S.M.K., Guzman E., Buccino A.P., Luna G., Cheng Z., Audouard M., Ranasinghe K.G., Kudo K., Nagarajan S.S., Tovar K.R., Petzold L.R., Hierlemann A., Hansma P.K. & Kosik K.S. Functional neuronal circuitry and oscillatory dynamics in human brain organoids. Nat Commun. 2022;13(1):4403.

53. Passaro A.P. & Stice S.L. Electrophysiological analysis of brain organoids: current approaches and advancements. Front Neurosci. 2020;14:622137.

54. Kobolak J., Teglasi A., Bellak T., Janstova Z., Molnar K., Zana M., Bock I., Laszlo L. & Dinnyes A. Human induced pluripotent stem cell-derived 3D-neurospheres are suitable for neurotoxicity screening. Cells. 2020;9(5).

