## Supplementary Figure 1 for "An optimized workflow to generate and characterize iPSC-derived motor neuron (MN) spheroids"

**This PDF file includes:**

Supplementary Figures 1-3

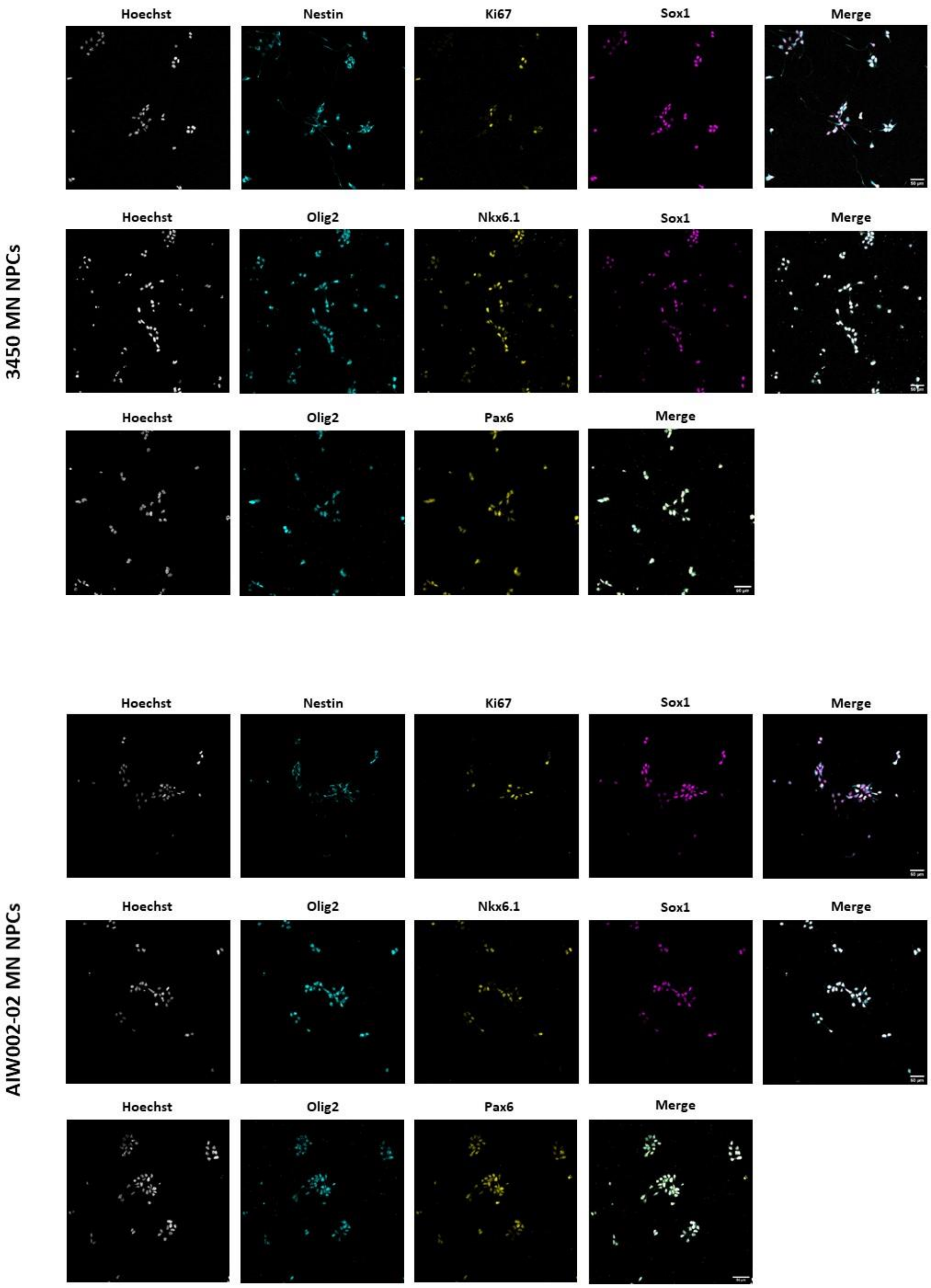


**Supplementary Figure 1. Motor neuron neural progenitor cell (MNPC) characterization by immunofluorescent staining.** MNPCs were positive for Sox1 and Nestin indicating their neural progenitor identity as well as Ki67, which demonstrates their proliferation capacity. Additionally, MNPCs are positive for the markers Pax6, Olig2 and NKx6.1 that indicate their successful specification towards motor neuron neural progenitor cells.
