## Supplementary Figure 2 for "An optimized workflow to generate and characterize iPSC-derived motor neuron (MN) spheroids"

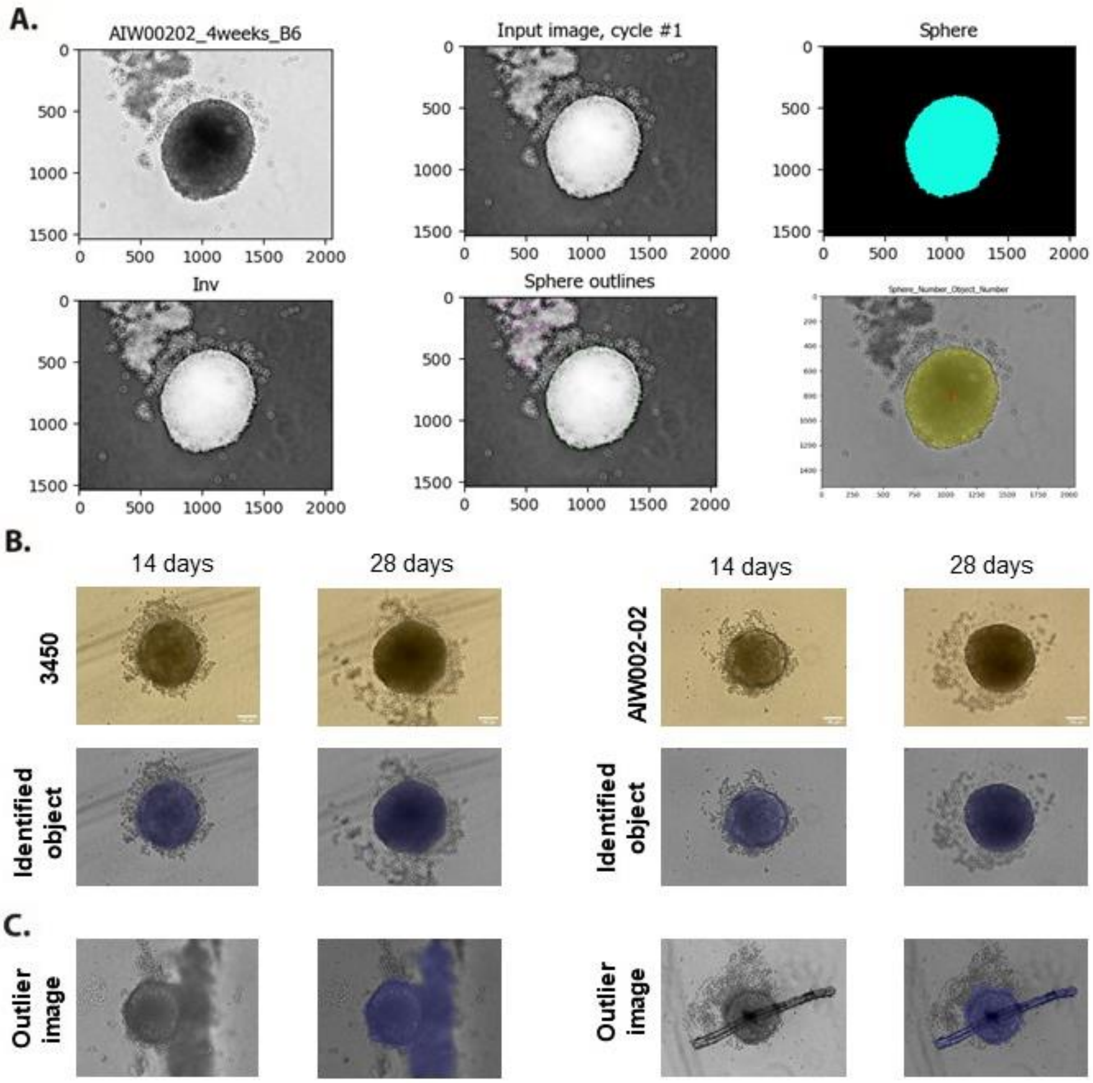


**Supplementary Figure 2. Cell Profiler macro to perform the size profiling of the MN spheroids. A)** A data set of pictures taken with a bright-field microscope is inserted into a Cell Profiler pipeline. **B)** The pipeline processes the image to identify a primary object that is overlaid with the original image. **C)** Images in which the macro performed poorly are considered outliers and removed from the analyses.
