## Supplementary Figure 3 for "An optimized workflow to generate and characterize iPSC-derived motor neuron (MN) spheroids"

**A.**

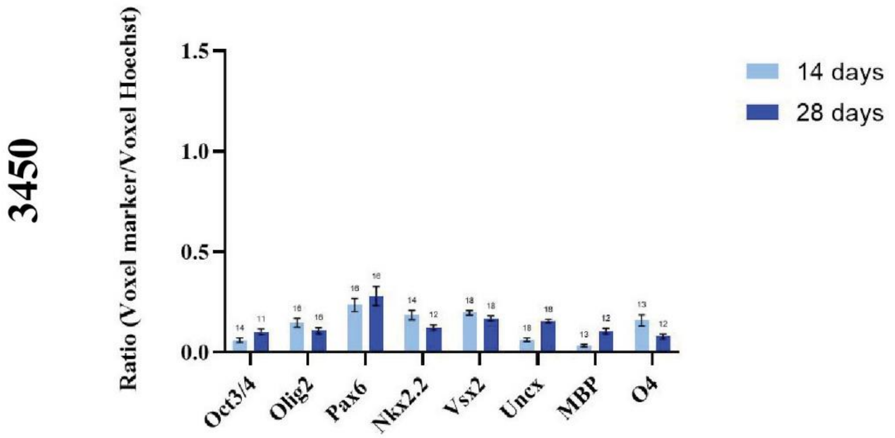


**B.**

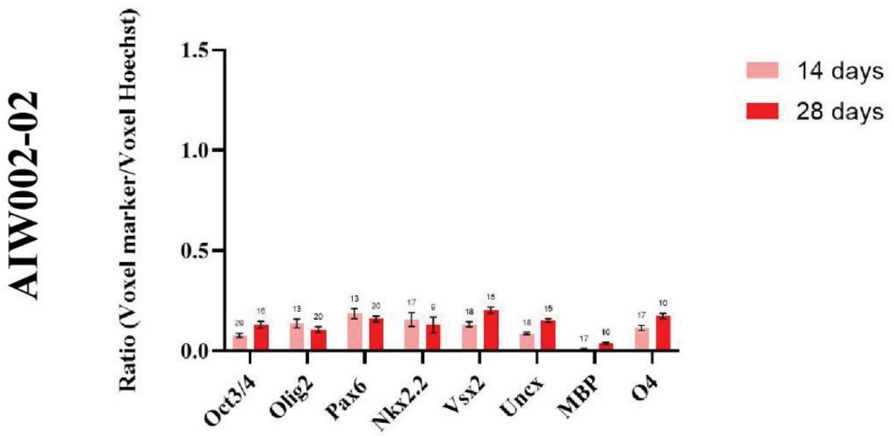


**Supplementary Figure 3. Image profiling of iPSC-derived MN spheroids to identify different cell types.** MN spheroids from **A)** 3450 and **B)** AIW002-02 control cell lines were stained for interneuron (Nkx2.2, Vsx2, Uncx) and oligodendrocyte markers (MBP, O4). The presence of these markers was quantified using an in-house MATLAB pipeline. Graph bars show the mean ± SEM; for each cell line, each batch of three batches of iPSC-derived MNPCs generated through independent differentiation processes was used to generate two MN spheroid batches. A minimum of 9 MN spheroids were required for quantification, and we ensured that at least 3 spheroids per MNPC batch where stained for each marker per cell line at each time point.
